# Anticancer pan-ErbB inhibitors reduce inflammation and tissue injury and exert broad-spectrum antiviral effects

**DOI:** 10.1101/2021.05.15.444128

**Authors:** Sirle Saul, Marwah Karim, Luca Ghita, Pei-Tzu Huang, Winston Chiu, Verónica Durán, Chieh-Wen Lo, Sathish Kumar, Nishank Bhalla, Pieter Leyssen, Farhang Alem, Niloufar A. Boghdeh, Do HN Tran, Courtney A. Cohen, Jacquelyn A. Brown, Kathleen E. Huie, Courtney Tindle, Mamdouh Sibai, Chengjin Ye, Ahmed Magdy Khalil, Luis Martinez-Sobrido, John M. Dye, Benjamin A. Pinsky, Pradipta Ghosh, Soumita Das, David E. Solow-Cordero, Jing Jin, John P. Wikswo, Dirk Jochmans, Johan Neyts, Steven De Jonghe, Aarthi Narayanan, Shirit Einav

**Author notes:** These authors contributed equally (Sirle Saul, Marwah Karim). These authors also contributed equally (Luca Ghita, Pei-Tzu Huang).

## Abstract

Targeting host factors exploited by multiple viruses could offer broad-spectrum solutions for pandemic preparedness. Seventeen candidates targeting diverse functions emerged in a screen of 4,413 compounds for SARS-CoV-2 inhibitors. We demonstrated that lapatinib and other approved inhibitors of the ErbB family receptor tyrosine kinases suppress replication of SARS-CoV-2, Venezuelan equine encephalitis virus (VEEV), and other emerging viruses with a high barrier to resistance. Lapatinib suppressed SARS-CoV-2 entry and later stages of the viral life cycle and showed synergistic effect with the direct-acting antiviral nirmatrelvir. We discovered that ErbB1, 2 and 4 bind SARS-CoV-2 S1 protein and regulate viral and ACE2 internalization, and they are required for VEEV infection. In human lung organoids, lapatinib protected from SARS-CoV-2-induced activation of ErbB-regulated pathways implicated in non-infectious lung injury, pro-inflammatory cytokine production, and epithelial barrier injury. Lapatinib suppressed VEEV replication, cytokine production and disruption of the blood-brain barrier integrity in microfluidic-based human neurovascular units, and reduced mortality in a lethal infection murine model. We validated lapatinib-mediated inhibition of ErbB activity as an important mechanism of antiviral action. These findings reveal regulation of viral replication, inflammation, and tissue injury via ErbBs and establish a proof-of-principle for a repurposed, ErbB-targeted approach to combat emerging viruses.

## Introduction

Acute emerging RNA viral infections can cause epidemics and pandemics and thus pose major threats to human health. Severe acute respiratory syndrome coronavirus 2 (SARS-CoV-2) has spread globally, causing largely asymptomatic or mild infections, yet progressing in some patients to severe Coronavirus Disease 2019 (COVID-19) manifesting with acute lung injury (ALI), acute respiratory distress syndrome (ARDS) and lung fibrosis (1, 2). Venezuelan Equine Encephalitis Virus (VEEV), an alphavirus naturally transmitted by mosquitoes, causes potentially lethal encephalitis associated with long-term neurological deficits in up to 14% of infected individuals (3). The most widespread mosquito-borne flavivirus, dengue (DENV), and the filoviruses Ebola (EBOV) and Marburg (MARV) are causative agents of outbreaks of potentially fatal hemorrhagic fever. Retaining stability and infectivity as an aerosol or droplets, VEEV and the filoviruses, respectively, are considered bioterrorism threats (3, 4). The current lack of effective intervention strategies against the majority of these and other emerging pathogens leave the global and military populations unprepared for future pandemics.

By targeting viral enzymes, the majority of approved antiviral strategies to date provide narrow-spectrum coverage. This approach has shown substantial utility in treating chronic viral infections, such as hepatitis C virus (HCV), and more recently, COVID-19. Nevertheless, the rapid rollout of direct-acting antivirals (DAAs) for COVID-19 treatment was enabled via either repurposing—remdesivir and molnupiravir, originally developed for Ebola virus disease and flu, respectively—or accelerated derivatization of existing SARS-CoV-1 main protease (Mpro) inhibitor in the case of nirmatrelvir. No such DAAs are available or in the pipeline for the majority of viral families, and since this approach to drug development is slow and expensive, it is not easily scalable to address the large unmet clinical need (5). Moreover, targeting viral factors by monotherapy often results in rapid emergence of drug resistance (5). Indeed, escape mutations conferring high-level resistance to remdesivir and nirmatrelvir have already been selected *in vitro* and identified in circulating SARS-CoV-2 strains (6, 7), and viral rebound, whose incidence is higher following paxlovid treatment, provides optimal conditions for the emergence of resistant mutants (8).

There is thus an unmet need for additional approaches, ideally targeting distinct mechanisms and increasing the barrier to resistance, to be used individually or in combination drug treatment for preventing the acute and long-term complications associated with viral infections and providing readiness for future outbreaks. Targeting host factors commonly required by multiple viral pathogens is an alternative antiviral approach that could provide broad-spectrum coverage while increasing the barrier to resistance (5, 9). The opportunity to repurpose existing drugs known to modulate specific host functions with favorable toxicity profiles is attractive, particularly for the treatment of emerging viral infections lacking any treatment.

To address these gaps, we conducted a high-throughput screen of existing compounds for agents that rescue mammalian cells from SARS-CoV-2-induced lethality. Among the hits were inhibitors of members of the epidermal growth factor receptor (ErbB) family of receptor tyrosine kinases, including lapatinib, an approved anticancer drug. Here, we reveal that ErbBs regulate both the life cycle and pathogenesis of SARS-CoV-2 and VEEV infections. Moreover, we provide support for the feasibility of repurposing pan-ErbB inhibitors as a candidate broad-spectrum antiviral, anti-inflammatory and tissue protective approach using *in vitro* and unique *ex vivo* models of multiple unrelated viral infections and a murine model of VEEV. Lastly, we characterize the mechanism of action of lapatinib and validate ErbBs as critical mediators of the antiviral effect.

## Results

### Pan-ErbB inhibitors emerge in a high-throughput screening (HTS) for compounds that counteract SARS-CoV-2-induced lethality

We assembled a collection of 4,413 bioactive investigational and FDA approved compounds derived from four commercially available libraries and a self-assembled set of 13 kinase inhibitors (**Figure 1A and Supplemental Figure 1A**). This collection was screened in two independent experiments for inhibition of lethality induced by SARS-CoV-2 (isolate: Belgium-GHB-03021) infection in Vero E6 cells constitutively expressing an enhanced green fluorescent protein (eGFP) via a high-throughput assay (10) (**Figure 1B**) that demonstrated robustness and specificity (**Supplemental text 1, Supplemental Figure 1B-D**). We set a percent fluorescent area of greater than 15 in at least one of the screens as the cutoff for positive hits (**Figure 1C and Supplemental Figure 1B-E**). Forty-two compounds, including nelfinavir and salinomycin previously demonstrating anti-SARS-CoV-2 activity (11), met this criterion. Eighteen of the 42 hits were prioritized based on PubChem data documenting lower promiscuity and toxicity or activity against other viruses (**Figure 1A**) and assessed for their effect on SARS-CoV-2 infection and cellular viability in Vero cells infected with a distinct viral isolate (2019-nCoV/USA-WA1/2020) via plaque and alamarBlue assays, respectively. In total, 17 hits demonstrated antiviral effect beyond toxicity, of which seven showed potent dose-dependent antiviral activity with EC_50_ (half-maximal effective concentration) <0.7 µM, CC_50_ (half-maximal cellular cytotoxicity) >20 µM, and selectivity indices (SI, CC_50_ to EC_50_ ratio) >20. These compounds target diverse cellular factors and functions (**Figure 1D-F and Supplemental Figure 2A**). Two of these hits were reported to target ErbBs: lapatinib and tyrphostin AG 879 (12). Inhibitors of NUMB-associated kinases (NAK), heat shock protein 90 (HSP90) and ion transport across cell membranes were also among the hits.

**Figure 1.**
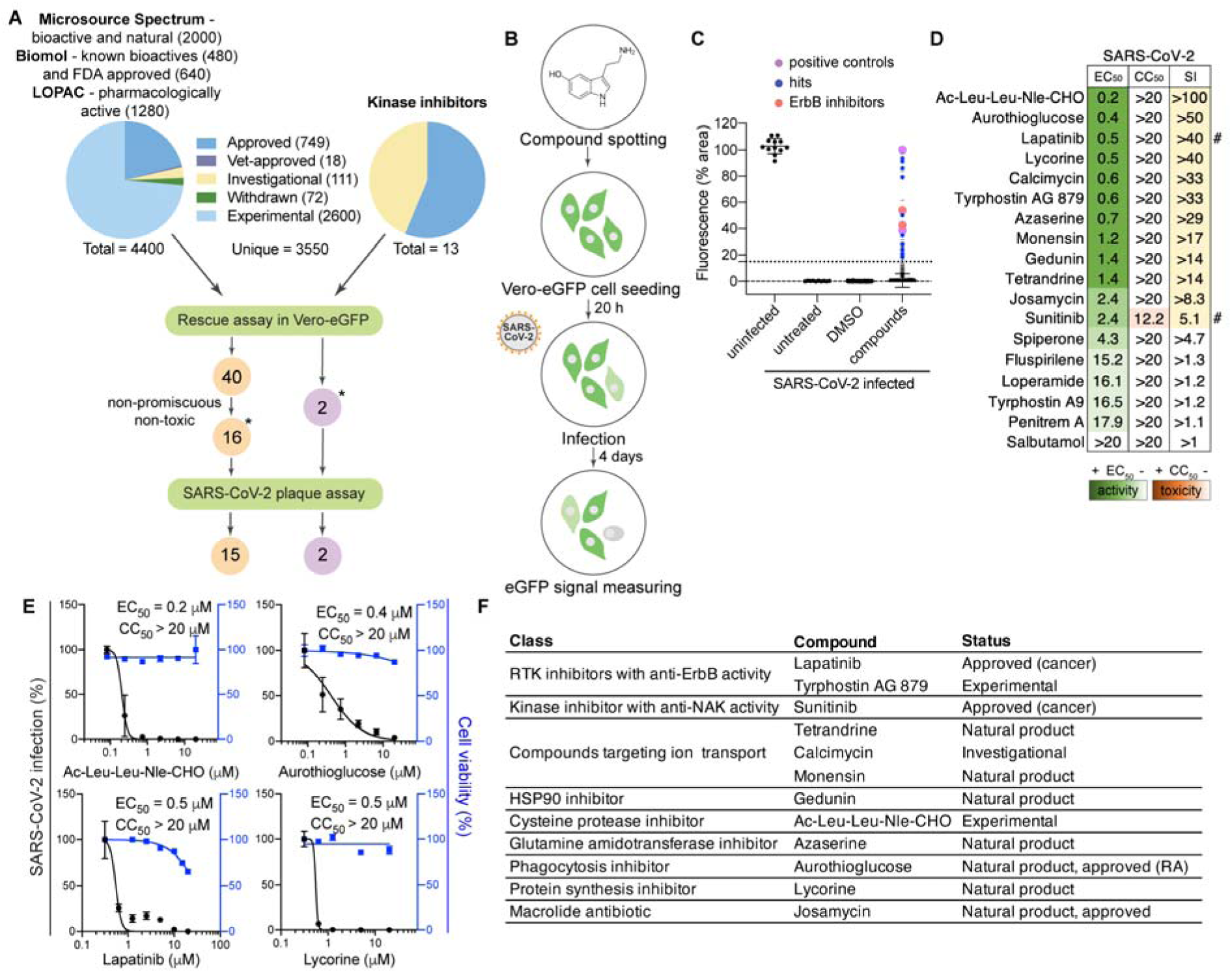
High-throughput screening (HTS) for compounds that counteract SARS-CoV-2-induced lethality and validation by plaque assays. **A**, A schematic of the composition of the screened libraries and screening and hit selection pipeline. **B**, HTS assay schematic. Compounds were pre-spotted in 384-well plates at a final concentration of 10 μM and incubated with Vero E6 cells constitutively expressing eGFP for 20 hours, followed by SARS-CoV-2 infection (Belgium-GHB-03021, MOI = 0.001). eGFP signal measured at 4 days post-infection was used as an indicator for survival from viral-induced lethality. **C**, Boxplots of the percentage of fluorescence area values combining the entire HTS data set (two independent experiments) split into the four indicated categories. The box horizontal lines indicate the first, second (median), and third quartiles. Outliers above a cutoff of 15% were defined as positive hits. Dots represent individual compounds and colors denote positive controls (purple), new hits (blue), and ErbB inhibitors (peach). **D**, Heat map of the EC_50_ and CC_50_ values of hits emerging in the HTS color-coded based on the antiviral activity measured by plaque assays (green) and toxicity measured by alamarBlue assays (orange) 24 hpi of Vero cells with SARS-CoV-2 (USA-WA1/2020 strain; MOI=0.05). Selectivity indices (SI) greater than 5 are depicted in yellow. # indicates compounds from the 13-kinase set. **E**, Representative dose-response curves of hits depicting SARS-CoV-2 infection (black) and cell viability (blue). Data are relative to DMSO. **F**, The 12 most promising hit compounds emerging in the HTS. Data in panels **D, E** are representative of 2 or more independent experiments. Individual experiments had three biological replicates. Shown are means ± SD. RA, Rheumatoid arthritis; RTK, receptor tyrosine kinase; NAK, NUMB-associated kinase. * in panel A denotes 18 hits screened for SARS-CoV-2, VEEV (TC-83) and DENV2.

### Lapatinib inhibits SARS-CoV-2 infection *in vitro* and *ex vivo* in human adult lung organoid (ALO)-derived monolayers and is highly synergistic with nirmatrelvir

Since lapatinib is an already approved, oral pan-ErbB inhibitor, we focused on defining its antiviral potential. Similarly to its effect in Vero cells (EC_50_=0.5 µM, CC_50_ >20 µM) (**Figure 1E**), in Calu-3 (human lung epithelial cells), lapatinib dose-dependently inhibited replication of SARS-CoV-2 (USA-WA1/2020 strain) as measured via plaque assay (EC_50_=0.7 µM), without apparent effect on cellular viability at the concentrations used as measured in the infected cells via alamarBlue assay (CC_50_ >20 µM) (**Figure 2A, B and 1E**). Likewise, lapatinib treatment dose-dependently suppressed infection of Calu-3 and Vero cells with replication-restricted pseudovirus bearing SARS-CoV-2 spike (S) protein (rVSV-SARS-CoV-2-S) as measured by luciferase assays (EC_50_=2.6-3.2 µM, CC_50_ >20 µM) (**Figure 2C, D**), suggesting that lapatinib inhibits viral entry. Moreover, lapatinib demonstrated a dose-dependent rescue of Vero-eGFP cells from SARS-CoV-2-induced lethality (**Figure 2E, F**).

**Figure 2.**
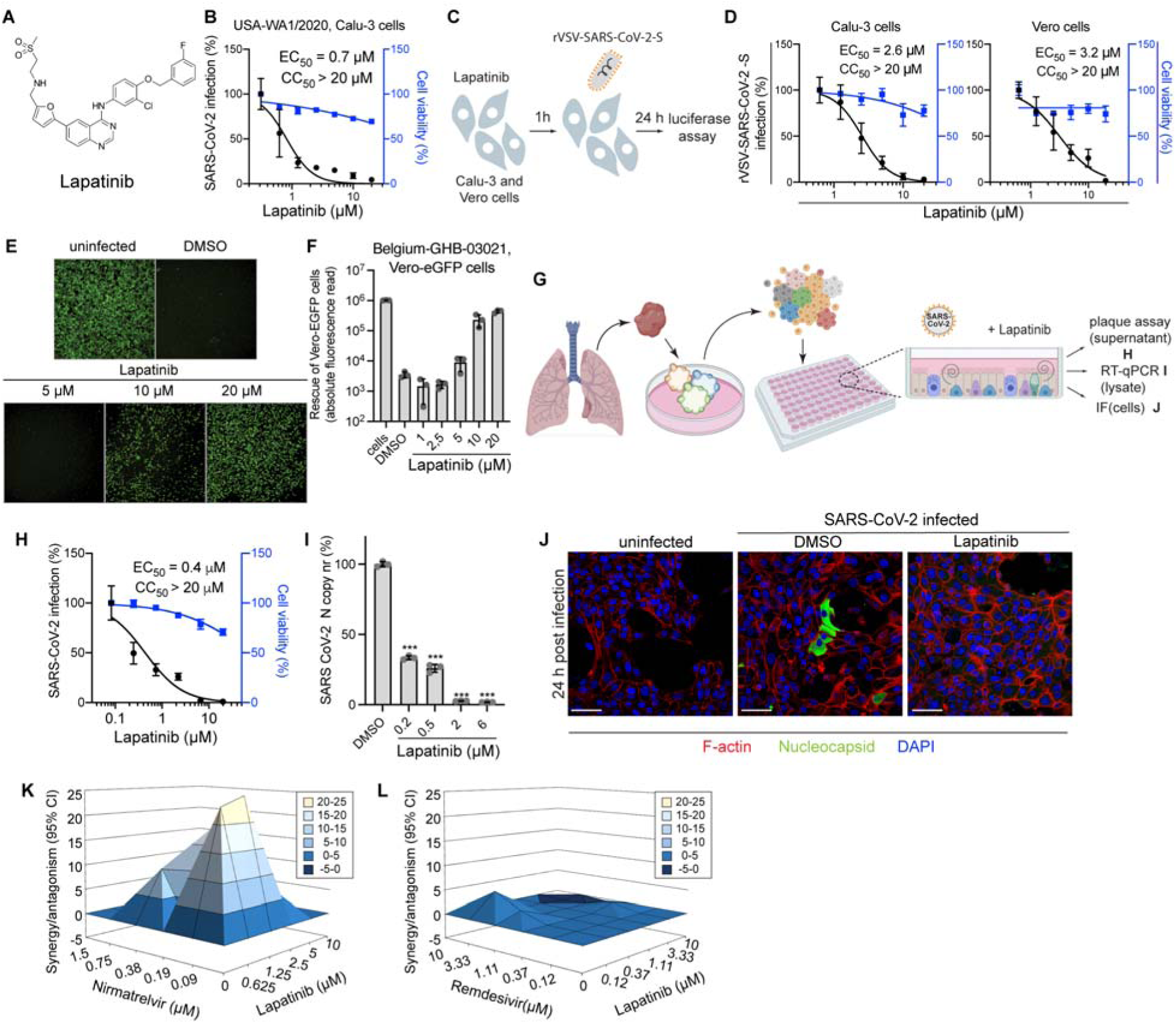
Lapatinib inhibits SARS-CoV-2 infection both *in vitro* and *ex vivo* and is synergistic with nirmatrelvir. **A,** Chemical structure of lapatinib. **B,** Dose response to lapatinib of SARS-CoV-2 infection (black, USA-WA1/2020 strain; MOI=0.05) and cell viability (blue) in Calu-3 cells measured via plaque and alamarBlue assays at 24 hpi, respectively. **C,** Schematic of the experiment shown in panel D. **D,** Dose response to lapatinib of rVSV-SARS-CoV-2-S infection (black) and cell viability (blue) in Calu-3 and Vero cells measured via luciferase and alamarBlue assays at 24 hpi, respectively. **E, F,** Dose-dependent graph (**F**) and corresponding florescence images (**E**) of Vero-eGFP cells rescued from SARS-CoV-2-induced lethality by lapatinib at 96 hpi (Belgium-GHB-03021 strain; MOI=0.05). **G,** Schematic of the ALO model and the experimental procedures. ALO-derived monolayers were infected with virulent SARS-CoV-2 (USA-WA1/2020 strain, MOI=1). **H,** Dose response to lapatinib of SARS-CoV-2 infection (black) and cell viability (blue) in ALO-derived monolayer supernatants measured via plaque and alamarBlue assays at 48 hpi, respectively. **I**, Dose response to lapatinib of SARS-CoV-2 nucleocapsid copy number in ALO-derived monolayer lysates measured by RT-qPCR assays at 48 hpi. **J,** Confocal IF microscopy images of F-actin (red), SARS-CoV-2 nucleocapsid (green) and DAPI (blue) in naïve and SARS-CoV-2-infected ALO-derived monolayers pre-treated with DMSO or 10 µM lapatinib 24 hpi. Representative merged images at 40x magnification are shown. Scale bars are 50 µm**. K, L** Synergy/antagonism of lapatinib and nirmatrelvir (**K**) or remdesivir (**L**) combination treatment on antiviral effect measured in Calu-3 cells infected with rSARS-CoV-2/Nluc (USA-WA1/2020 strain; MOI = 0.05) at 24 hpi via Nluc assays. Data represent differential surface analysis at the 95% confidence interval (CI), analyzed via the MacSynergy II program. Data are representative of two independent experiments with three replicates each. Data in **B, D, H** and **I** are relative to DMSO. Means ± SD are shown. \*\*\**P* < 0.001 by 1-way ANOVA followed by Dunnett’s multiple comparisons test. PFU, plaque-forming units.

To study the effect of lapatinib treatment on SARS-CoV-2 infection in a more biologically relevant model, we used a validated human adult lung organoid (ALO)-derived monolayer model. Generated from adult stem cells isolated from lung tissue, these organoid-derived monolayers contain both proximal airway cells, critical for sustained viral infection, and distal alveolar cells, required for mounting the overzealous host immune response in fatal COVID-19 (13) (**Figure 2G**). Viral replication measured by plaque assays in culture supernatant and nucleocapsid transcript expression measured by RT-qPCR in ALO-derived monolayer lysates both peaked at 48 hours following SARS-CoV-2 infection (**Supplemental Figure 2B, C**) and were effectively and dose-dependently suppressed by lapatinib, with EC_50_ values of 0.4 µM and <0.2 µM, respectively, and CC_50_ > 20 µM (**Figure 2H, I**). Confocal immunofluorescence (IF) analysis revealed a near-complete disappearance of SARS-CoV-2 nucleocapsid staining in ALO-derived monolayers treated with 10 µM of lapatinib relative to DMSO controls (**Figure 2J and Supplemental Figure 2D**).

To determine the utility of lapatinib in combination treatment, we measured the anti-SARS-CoV-2 activity of combinations of lapatinib with clinically used DAAs. Treatment with lapatinib-nirmatrelvir combinations exhibited synergistic inhibition of SARS-CoV-2 infection as measured via luciferase assay with a synergy volume of 91.42 μM^2^% (within a range that is considered moderate and probably important *in vivo* (14)) and antagonism volume of 0 μM^2^% at the 95% confidence interval (MacSynergy (14)) (**Figure 2K**). In contrast, lapatinib-remdesivir combinations were additive (**Figure 2L**). No synergistic toxicity was measured with these combinations via alamarBlue assay (**Supplemental Figure 2E, F)**.

These results point to lapatinib as a potent anti-SARS-CoV-2 inhibitor with potential utility in combination drug treatment with paxlovid.

### Lapatinib has a broad-spectrum antiviral activity and a high genetic barrier to resistance

Next, we studied the broad-spectrum potential of lapatinib and the other 17 hits emerging from the HTS (**Supplemental text 2, Supplemental Figure 3A, B**). Lapatinib dose-dependently inhibited alphavirus replication of both vaccine (TC-83) and wild type (WT) (Trinidad donkey, TrD) VEEV by plaque assays in human astrocytes (U-87 MG) cells with EC_50_ values of 1.2 μM and 0.8 μM respectively and CC_50_ >20 μM (**Figure 3A, B**). Similarly, lapatinib dose-dependently inhibited the replication of the flavivirus DENV2 (EC_50_=1.8 μM) via plaque assays, and the filoviruses EBOV (EC_50_=2.5 μM) and MARV (EC_50_=1.9 μM) via microneutralization assays in human hepatoma (Huh7) cells, albeit lower CC_50_ values were measured in infected Huh7 cells (10.2-10.5 μM) relative to the other cell lines (**Supplemental Figure 4A-C**).

**Figure 3.**
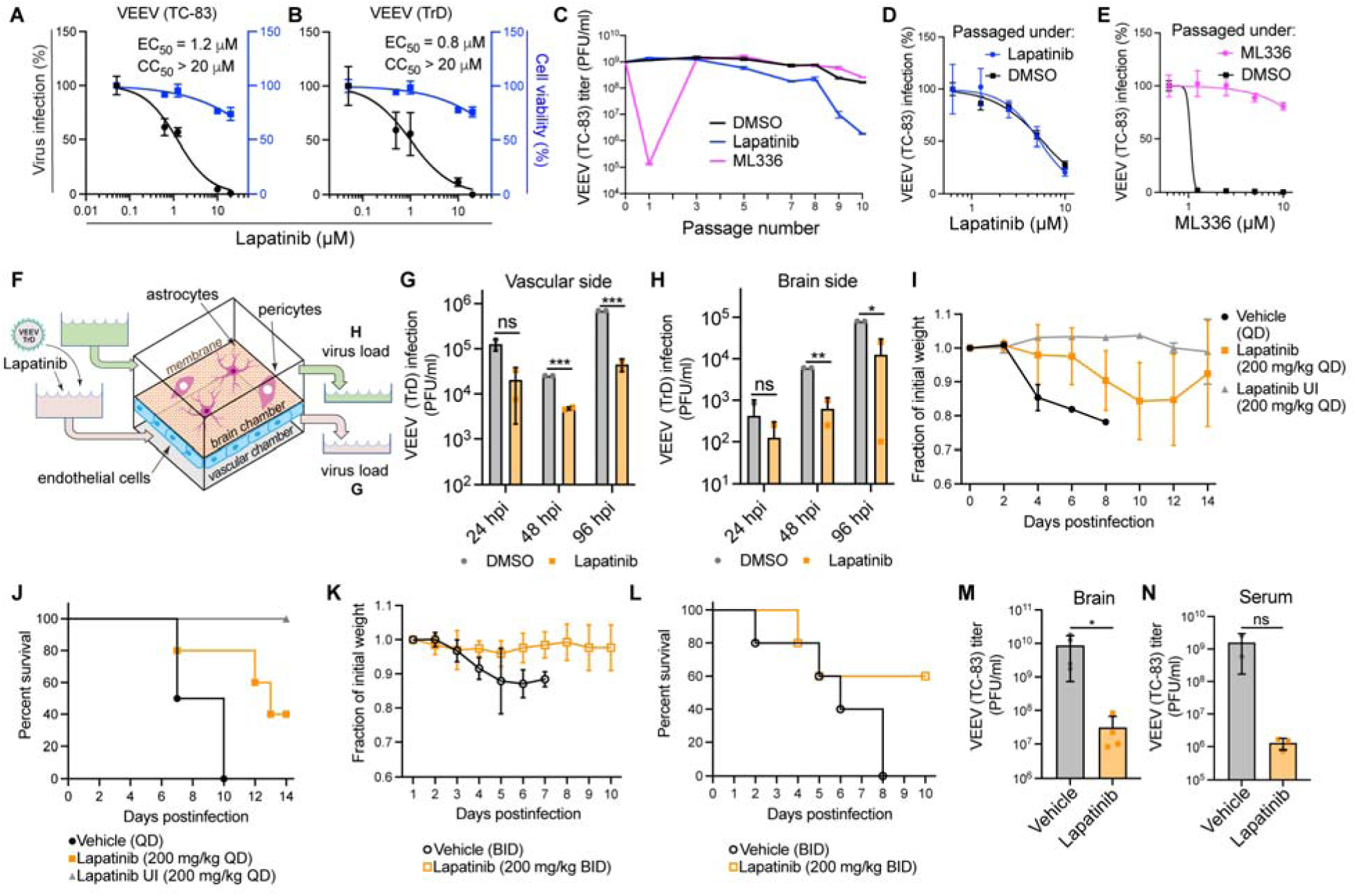
Lapatinib is a potent broad-spectrum antiviral with a high genetic barrier to resistance and is protective in human neurovascular units (gNVU) and murine model of VEEV. **A, B** Dose response to lapatinib of infection with vaccine (TC-83) (**A**) and WT (Trinidad donkey (TrD)) (**B**) VEEV strains (MOI=0.1) in U-87 MG cells via plaque and alamarBlue assays at 24 hpi, respectively. **C,** VEEV (TC-83) was used to infect U-87 MG cells (MOI=0.1) and was then passaged every 24 hours by inoculation of naive U-87 MG cells with equal volumes of viral supernatants under DMSO treatment or selection with lapatinib or ML336 (VEEV nsP2 inhibitor) increasing from 2.5 to 15 μM over 10 passages. Viral titers were measured by plaque assays. **D, E** Dose response to lapatinib (**D**) and ML336 (**E**) of VEEV (TC-83) harvested after 10 passages in U-87 MG cells in the presence of lapatinib (**D**) and ML336 (**E**), via luciferase assays. **F**, Schematic of the gNVU. **G, H** Viral load in longitudinal samples collected from the vascular (**G**) and brain (**H**) sides of the gNVU following infection with VEEV (TrD) and treatment with lapatinib or DMSO. **I-L** Weight loss (**I, K**) and mortality (**J, L**) of VEEV (TC-83)-infected C3H/HeN mice treated once (I, J) or twice (K, L) daily for 14 (I, J) or 10 (K, L) days with vehicle or lapatinib (200mg/kg) (*n*=5 per treatment group). **M, N** Titers of VEEV (TC-83) in brain (**M**) and serum (**N**) samples from mice treated twice daily (*n*=3 per treatment group). \**P* < 0.05, \*\**P* < 0.01, \*\*\**P* < 0.001 by unpaired t-test (G, H) or nonparametric Mann-Whitney test (M, N). Data in A, B, D, E are relative to DMSO. Data in panels A, B, D, E are representative of 2 independent experiments. Means±SD of results of representative experiments conducted with three (A-E) or two (G, H) replicates are shown. QD, once daily; BID, twice daily; UI, uninfected; ns, non-significant.

To determine whether viruses can escape treatment with lapatinib, VEEV (TC-83) was passaged in U-87 MG cells in the presence of lapatinib or the VEEV nonstructural protein 2 (nsP2) inhibitor ML336 (15) at increasing concentrations corresponding to values between the EC_50_ and EC_90_, and viral titers were measured in culture supernatants by plaque assays. By passage 3, VEEV overcame inhibition by ML336. In contrast, VEEV remained suppressed for 10 passages under lapatinib treatment without phenotypic resistance (**Figure 3C**). Moreover, virus obtained from culture supernatants at passage 10 under lapatinib or DMSO treatment remained susceptible to lapatinib (**Figure 3D**). Conversely, virus obtained at passage 10 under ML336 treatment lost its susceptibility to ML336, with the emergence of a previously characterized resistance mutation in nsP2 (Y102C in VEEV TC-83), whereas virus obtained at the same passage under DMSO treatment remained susceptible to ML336 (**Figure 3E**).

These results point to lapatinib as a potential broad-spectrum antiviral agent with a higher relative barrier to resistance than a DAA, and support that lapatinib suppresses viral infection by targeting a cellular function.

### Lapatinib suppresses VEEV replication in an organ-on-a chip human neurovascular unit (NVU) model and protects mice from VEEV challenge

Since disruption of the blood-brain barrier (BBB) substantially contributes to encephalitic outcomes in the context of VEEV infection (16), we studied the effect of lapatinib treatment on the BBB integrity following infection with VEEV in a recently validated gravity-flow NVU (gNVU) model (17, 18). The gNVU—composed of human primary brain endothelial cells on one side of a membrane and astrocytes and pericytes on the other so as to establish independent brain and vascular chambers—recreates the dynamic of multicellular BBB microenvironment (18). Lapatinib- or DMSO-containing culture medium was perfused an hour prior to introduction of TrD-containing medium into the vascular inlet of the gNVU (**Figure 3F**). Lapatinib treatment (5 µM) suppressed VEEV (TrD) replication in both the vascular and brain sides of the gNVU, as measured by plaque assays in perfused media at 24, 48 and 96 hours postinfection (hpi) (**Figure 3G, H**).

To further address the therapeutic potential of lapatinib as an antiviral agent, we tested its application in a murine model of VEEV (TC-83) under BSL2 conditions. C3H/HeN mice were infected intranasally with a lethal infectious dose of VEEV (TC-83) inoculum (5×10^6^ PFU). Once-daily (QD) treatment with 200 mg/kg lapatinib or vehicle alone via oral gavage was initiated at 12 hours prior to inoculation. The dose tested was lower than both, the approved human dose, as calculated based on the body surface area per the FDA’s guidelines (19), and the maximum tolerated dose (MTD) in mice (20, 21) and confirmed to be nontoxic in our VEEV model. The animals were monitored twice daily and were euthanized when moribund. During a 14-day drug treatment, we observed a significant reduction in morbidity and mortality of infected animals relative to vehicle controls (**Figure 3I, J**). Specifically, whereas 100% of vehicle-treated mice succumbed to infection by day 10 postinfection, lapatinib treatment protected 80% of the mice by day 10, and 40% of the mice by day 14.

Since lapatinib’s half-life is shorter in mice than humans, we administered lapatinib (200 mg/kg) twice daily for 8 days in uninfected C57BL/6 mice, observing good tolerability and plasma concentrations exceeding the EC_50_ deduced from our *in vitro* data by 6-43 folds (**Supplemental Figure 4D**). Twice-daily (200mg/kg BID) 10-day lapatinib treatment in the VEEV (TC-83) murine model protected 60% of the mice, whereas 100% of vehicle-treated mice succumbed to infection by day 8 postinfection (**Figure 3K, L**).

To determine whether the improved disease outcome is associated with reduction in viral load, we assessed viral burden in serum and brain. We measured a 3-log reduction of the infectious virus load by plaque assays in both the serum and brain in mice treated twice daily with lapatinib relative to vehicle controls (**Figure 3M, N**).

Together, these results demonstrate therapeutic potential of lapatinib in biologically relevant models against infections with at least two unrelated emerging RNA viruses, SARS-CoV-2 and VEEV.

### ErbBs are essential for SARS-CoV-2 and VEEV infections

The ErbB family is composed of four members (ErbB1-4), of which three, ErbB1, 2, and 4, are catalytically active (22). Lapatinib’s cancer targets are ErbB1 (EGFR) (IC_50_=5.3 nM) and ErbB2 (HER2) (IC_50_=35 nM)(23), yet it was shown to bind the ATP binding site of ErbB4 in a comparable manner to its interaction with ErbB1 and ErbB2 (24). Indeed, we measured an IC_50_ of 28 nM of lapatinib on ErbB4 in a cell-free assay and confirmed its anti-ErbB2 activity (**Supplemental Figure 5A**). Beyond ErbBs, lapatinib’s kinome (ID:20107) reveals potent binding to RAF1, STK10, RIPK2, and MAP2K5, with an overall excellent selectivity for ErbBs by Kd measurements (ID:20155). To define the molecular targets mediating the observed antiviral effect of lapatinib, we studied the effects of siRNA-mediated depletion of these seven kinases in Vero E6 cells infected with WT SARS-CoV-2 via plaque assay (**Figure 4A, B**). ErbB depletion suppressed SARS-CoV-2 replication by ∼50% relative to a non-targeting (siNT) control. A similar phenotype was observed in Vero cells infected with pseudovirus (rVSV-SARS-CoV-2-S), revealing a role of ErbBs in viral entry, a stage of the SARS-CoV-2 life cycle that is inhibited by lapatinib (**Supplemental Figure 5B, C**). In Calu-3 cells, depletion of ErbBs by the same siRNA pools suppressed WT SARS-CoV-2 infection by 97-98% relative to siNT as measured via plaque assays (**Figure 4A, C**). Similarly, silencing ErbBs expression in U-87 MG cells by these siRNAs resulted in 1- to over 2-log suppression of TrD infection and 30-70% suppression of VEEV (TC-83) infection relative to siNT, as measured via plaque assays (**Figure 4A, D, Supplemental Figure 5D, E**). ErbB depletion by the siRNA pools did not impact cell viability (**Figure 4B-D, Supplemental Figure 5C, E**), and its efficiency was confirmed by Western blot and RT-qPCR analysis (**Figures 4E, F and Supplemental Figure 5F**). STK10 depletion inhibited both SARS-CoV-2 and TrD infections, whereas RIPK2 and RAF1 depletion suppressed only TrD infection (**Figures 4B, D**).

**Figure 4.**
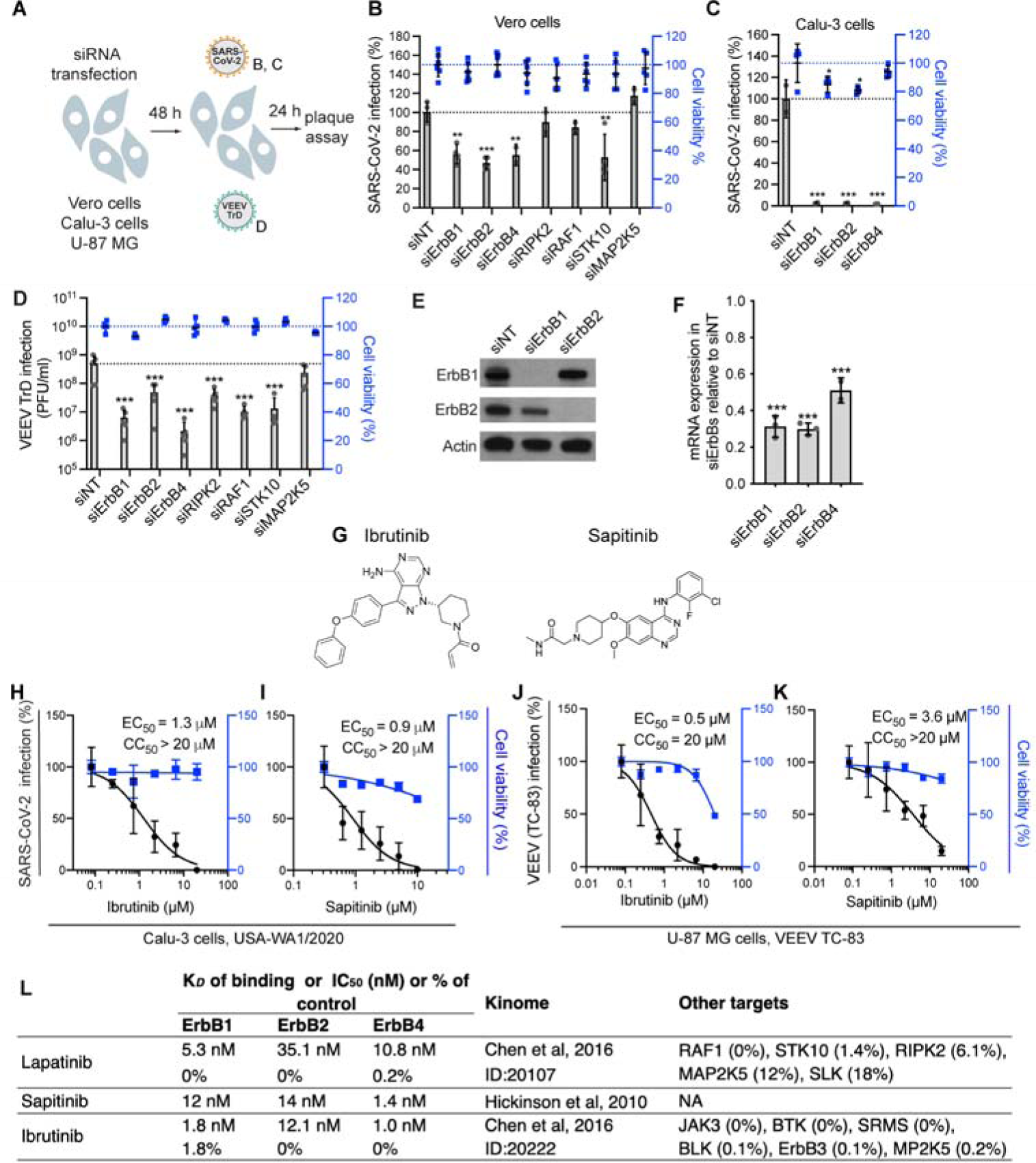
ErbBs are essential for SARS-CoV-2 and VEEV (TC-83) infections. **A**, Schematic of the experiment shown in panels B, C and D. **B, C,** Percentage of infection by plaque assays (grey) and cell viability by alamarBlue assays (blue) measured at 24 hpi of Vero (**B**) and Calu-3 cells (**C**) transfected with the indicated siRNA pools with WT SARS-CoV-2. **D**, Viral titers (grey) and cell viability (blue) measured at 24 hpi of U-87 MG cells transfected with the indicated siRNA pools with VEEV (TrD). **E, F**, Confirmation of siRNA-mediated gene expression knockdown in Vero cells at 48 hours post-transfection by Western blot (**E**) and RT-qPCR (**F**). Shown is gene expression normalized to GAPDH and expressed relative to the respective gene level in the non-targeting (siNT) control. Notably, two anti-ErbB4 antibodies detected no signal of endogenous protein in Vero cells. **G,** Chemical structures of ibrutinib and sapitinib. **H-K** Dose response to ibrutinib (**H, J**) and sapitinib (**I, K**) of SARS-CoV-2 (black, USA-WA1/2020 strain; MOI=0.05) (H, I) and VEEV (TC-83) (J, K) infection by plaque assays and cell viability (blue) by alamarBlue assays at 24 hpi of Calu-3 cells (H, I) or U-87 MG (J, K) cells. **L,** Binding affinity (K*_D_*), enzymatic activity (IC_50_) or percent binding of control (% control) of the indicated kinase inhibitors on the three catalytic ErbBs, the source of kinome data, and other targets these compounds bind and/or inhibit. Data in all panels are representative of two or more independent experiments. Individual experiments had three biological replicates, means ± SD are shown. Data are relative to DMSO (**H-K**) or siNT **(B, C, D, F**). \*\**P* < 0.01, \*\*\**P* < 0.001 by one-way ANOVA followed by Dunnett’s multiple comparisons test.

To further probe the requirement for ErbBs in SARS-CoV-2 and VEEV infections, we evaluated the antiviral effect of two chemically distinct compounds: ibrutinib, an approved anticancer Bruton’s tyrosine kinase (BTK) inhibitor, and sapitinib (investigational), both with potent pan-ErbB activity (23, 25) (**Figure 4G, Supplemental Figure 5G**). These compounds suppressed SARS-CoV-2 and VEEV (TC-83) infections, with EC_50_ values at sub to low micromolar range and CC_50_≥20 µM (**Figure 4H-K**).

These findings provide genetic and pharmacological evidence that ErbBs are required for SARS-CoV-2 and VEEV infections, thereby validating them as druggable antiviral targets.

### Mechanisms underlying the antiviral effects of lapatinib

#### ErbBs bind the viral spike S1 subunit, and their inhibition suppresses SARS-CoV-2 internalization

To better understand the mechanism of action underlying the anti-SARS-CoV-2 activity of lapatinib, we first probed the steps of the viral life cycle inhibited by lapatinib via time-of-addition experiments. Lapatinib was added to Calu-3 cells upon infection or at 2, 5 or 8 hpi with WT SARS-CoV-2 (**Figure 5A**). Cell culture supernatants were harvested at 10 hpi, which represented a single cycle of viral replication in Calu-3 cells, and infectious viral titers were quantified by plaque assays. Lapatinib treatment initiated upon infection onset and maintained throughout the 10-hour experiment (0 to 10) suppressed viral infection by 98% (**Figure 5B**). Lapatinib treatment during the first 2 hours of infection only (0 to 2) suppressed viral infection by 75%, confirming an effect on entry of WT SARS-CoV-2 (beyond rVSV-SARS-CoV-2-S, **Figure 2D**). Nevertheless, the addition of lapatinib at 2, 5 and 8 hpi suppressed viral infection by 99%, 84% and 57%, respectively, indicating inhibition also at post-entry stages (**Figure 5B**).

**Figure 5.**
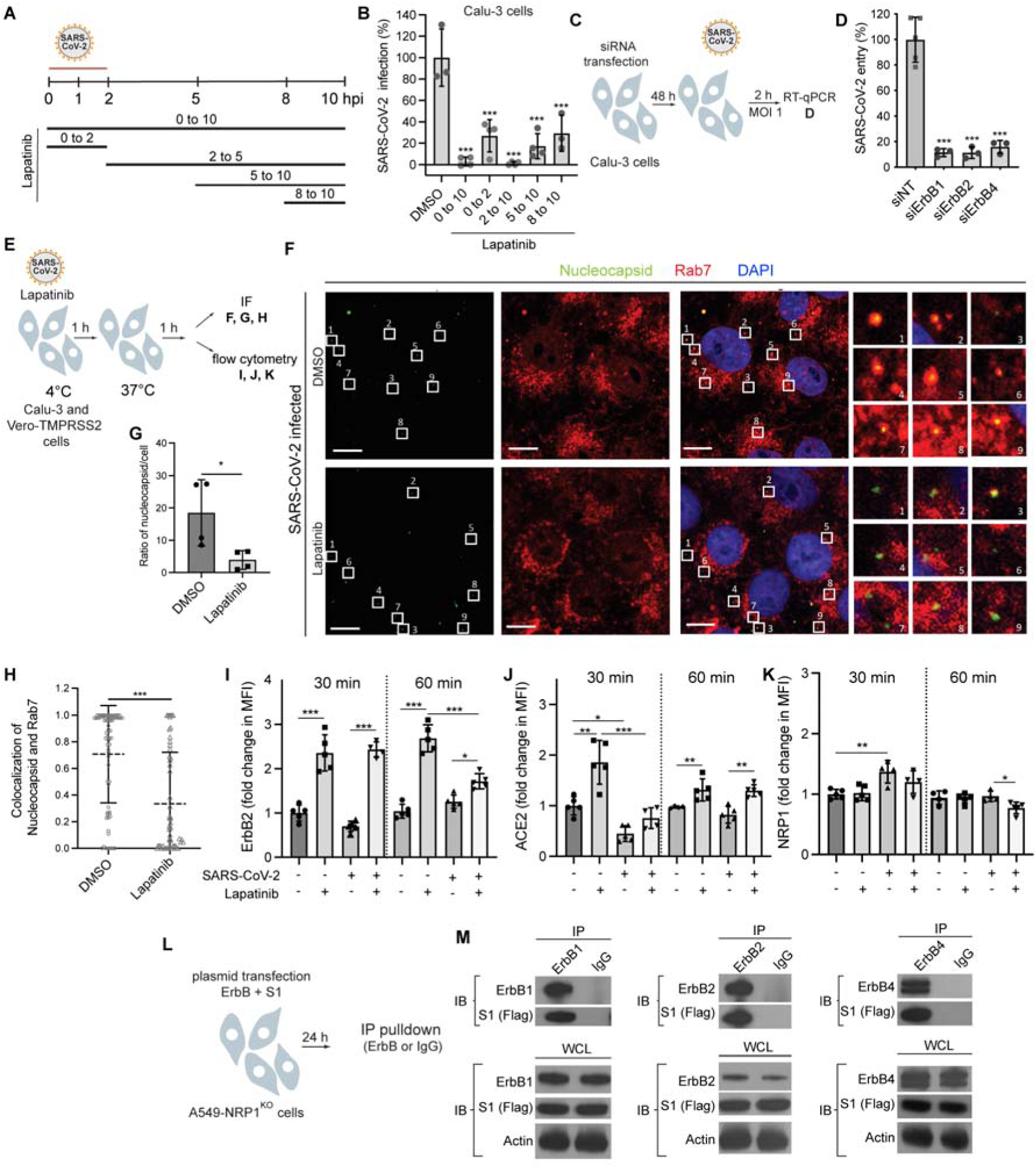
ErbBs bind the viral spike S1 subunit, and their inhibition suppresses SARS-CoV-2 and ACE2 internalization. **A**, Schematic of the time-of-addition experiment shown in panel B**. B,** Calu-3 cells were infected with SARS-CoV-2 (MOI=1). At the indicated time points, 10 µM lapatinib or DMSO were added to the infected cells. Supernatants were collected 10 hpi and infectious virus titers were measured by plaque assay. Values are shown relative to DMSO control. **C**, Schematic of the experiment shown in panel D. **D,** WT SARS-CoV-2 entry at 2 hpi of Calu-3 cells (MOI=1) depleted of the indicated ErbBs measured by RT-qPCR. **E**, Schematic of the experiments shown in panels F-K. **F-H** Quantitative IF analysis of SARS-CoV-2 internalization. Vero-TMPRSS2 cells were pretreated with lapatinib (10 µM) or DMSO and infected with SARS-CoV-2 (MOI=1) at 4°C for 1 hour followed by temperature shift to 37°C. At 1 hpi, cells were fixed and labeled with nucleocapsid (green) and Rab7 (red) antibodies. 6x zoom ins are shown. Scale bars are 10 µm (**F**). **G,** Ratio of viral particles per cell in DMSO- and lapatinib-treated cells. **H**, Scatter plots of colocalization of nucleocapsid and Rab7 quantified by Manders’ coefficient. Each dot represents an individual viral particle; horizontal lines indicate the means ± standard deviations (error bars) (*n*=71 and 53 in DMSO- and lapatinib-treated cells, respectively). **I-K,** Flow cytometry data of cell surface expression levels of ErbB2 (**I**), ACE2 (**J**) and NRP1 (**K**) at 30 and 60 min after temperature shift to 37°C in uninfected and SARS-CoV-2-infected Calu-3 cells treated with lapatinib or DMSO. The fold change in mean fluorescence intensity (MFI) is relative to 30 min uninfected DMSO treated cells. **L,** Schematic of the experiment shown in panel M. **M**, A549-NRP1^KO^ cells were cotransfected with plasmids expressing S1-Flag and ErbBs, followed by immunoprecipitation (IP) using anti-ErbB1, ErbB2, ErbB4 or IgG antibodies and protein G Dynabeads. Representative Western blots of eluates and whole-cell lysates (WCL) are shown. Data in all panels are representative of two or more independent experiments. Individual experiments had three biological replicates, means ± SD are shown (**B, D, G-K**). Data are relative to DMSO (**B, G-K**) or to siNT (**D**). \*\**P* < 0.01, \*\*\**P* < 0.001 by one-way ANOVA followed by Dunnett’s multiple comparisons test. Ns, non-significant.

To genetically confirm the role of ErbBs in the entry of WT SARS-CoV-2 (beyond rVSV-SARS-CoV-2-S, **Supplemental Figure 5B, C**), Calu-3 cells depleted for the individual ErbBs by the corresponding siRNAs were infected with high-inoculum virus followed by quantification of intracellular viral RNA at 2 hpi by RT-qPCR. siErbB1, siErbB2 and siErbB4 suppressed SARS-CoV-2 entry by 84-89% relative to siNT (**Figure 5C, D**).

To distinguish between viral binding and post-binding events, rVSV-SARS-CoV-2-S was incubated with Vero cells for 2 hours at 4°C in the presence or absence of lapatinib or DMSO before initiation of infection by shifting the temperature to 37°C (**Supplemental Figure 6A**). Lapatinib had comparable effects on rVSV-SARS-CoV-2-S infection when added upon or following binding of the virus to target cells with no apparent effect on cell viability (**Supplemental Figure 6B**), providing evidence for suppression at a post-binding step.

We next monitored the effect of lapatinib on single SARS-CoV-2 particle internalization in TMPRSS2-expressing Vero E6 cells fixed at 1 hpi and temperature shift to 37°C, labeled for viral particles and late endosomes with antibodies targeting the nucleocapsid and Rab7, respectively, and imaged by confocal microscopy. Quantitative IF analysis revealed that lapatinib treatment significantly reduced both the ratio of nucleocapsid puncta per cell and the colocalization of over 50 randomly chosen SARS-CoV-2 particles in each category to late endosomes, a cellular compartment into which SARS-CoV-2 internalizes (26), with mean Manders’ coefficients of 0.33 vs. 0.71 in DMSO-treated cells (**Figure 5E-H**). The majority of viral particles had an equivalent size based on fluorescence emissions, suggesting that single viral particles were imaged.

We tested the hypothesis that lapatinib alters the internalization of the SARS-CoV-2 receptor ACE2 and co-receptor neuropilin-1 (NRP1) (27, 28). The extracellular expression levels of ErbB2 and these cell surface receptors were measured by flow cytometry analysis of Calu-3 cells pretreated with lapatinib or DMSO, infected with SARS-CoV-2 (or mock) at 4°C, and extracellular staining at 30 and 60 min following a temperature shift to 37°C (**Figure 5E, I-K, Supplemental Figure 6C, D**). In uninfected cells, lapatinib treatment caused over 2-fold increase in the cell surface level of ErbB2 at both time points (**Figure 5I**), in agreement with its reported effect on dimerization and internalization of ErbB receptor complexes (beyond phosphorylation)(29). Interestingly, lapatinib had a similar effect, most prominently at 30 min post-infection, on the surface level of ACE2, but not NRP1 (**Figure 5J, K**). SARS-CoV-2 infection decreased ACE2 surface level at 30 min following temperature shift, suggesting that ACE2 internalization, previously shown to be caused by S1-ACE2 binding (30, 31), is induced during SARS-CoV-2 entry (**Figure 5J**). A trend towards reduced ErbB2 levels, albeit statistically nonsignificant, was also measured at the early time point post-infection, whereas the level of cell surface NRP1 was increased upon infection (**Figure 5I, K**). Lapatinib treatment reversed the effect of SARS-CoV-2 infection on the surface level of ACE2 (**Figure 5J**) and increased the surface level of ErB2, but not NRP1, in infected cells relative to DMSO (**Figure 5I, K**), suggesting that the internalization of ACE2 but not NRP1 may be regulated by ErbBs.

To determine whether ErbBs interact with the receptor binding domain of SARS-CoV-2 spike protein S1, we performed co-immunoprecipitation (IP) assays. Since S1 was shown to bind ACE2 and NRP1, we studied potential interactions of S1 with ErbBs in A549-NRP1^KO^ cells that are intrinsically deficient of ACE2 and deleted for NRP1 by CRISPR/Cas-9. A549-NRP1^KO^ cells were co-transfected with plasmids expressing Flag-tagged S1 and the individual ErbBs (**Figure 5L**). Anti-ErbB1, 2 and 4 antibodies effectively pulled down the respective ErbB (∼180 kDa), with which a ∼100 kDa protein corresponding to S1 was co-immunoprecipitated (**Figure 5M**). No background signal was demonstrated with control immunoglobulin G (IgG), indicating specificity of the observed IP.

Together, these results provide evidence that ErbBs regulate SARS-CoV-2 internalization and viral-induced ACE2 internalization, and that their direct, ACE2- and NRP1-independent, binding to the viral S1 subunit may be implicated in this process.

### ErbBs are the molecular targets mediating the antiviral effect of lapatinib

To determine whether lapatinib exerts its antiviral effect by inhibiting phosphorylation of ErbBs, lysates derived from SARS-CoV-2-infected Calu-3 cells treated with lapatinib or DMSO were subjected to quantitative Western blot analysis of phospho-ErbB to total ErbB ratios. SARS-CoV-2 infection induced mild phosphorylation of ErbB1 and ErbB2 in these cells. Lapatinib treatment dose-dependently suppressed the ratio of phosphorylated to total ErbB1, 2, and 4 levels at 24 hpi, with EC_50_ values lower than 0.1 µM, which correlated with reduced expression of the SARS-CoV-2 nucleocapsid protein (**Figure 6A, B**). Analogous suppression of ErbB phosphorylation was measured in Calu-3 cells at 1.5 hpi and in ALO monolayers at 1.5 and 24 hpi (**Supplemental Figure 7A-C**). These results provide evidence that drug exposure and the antiviral effect of lapatinib are correlated with functional inhibition of ErbBs’ activity.

**Figure 6.**
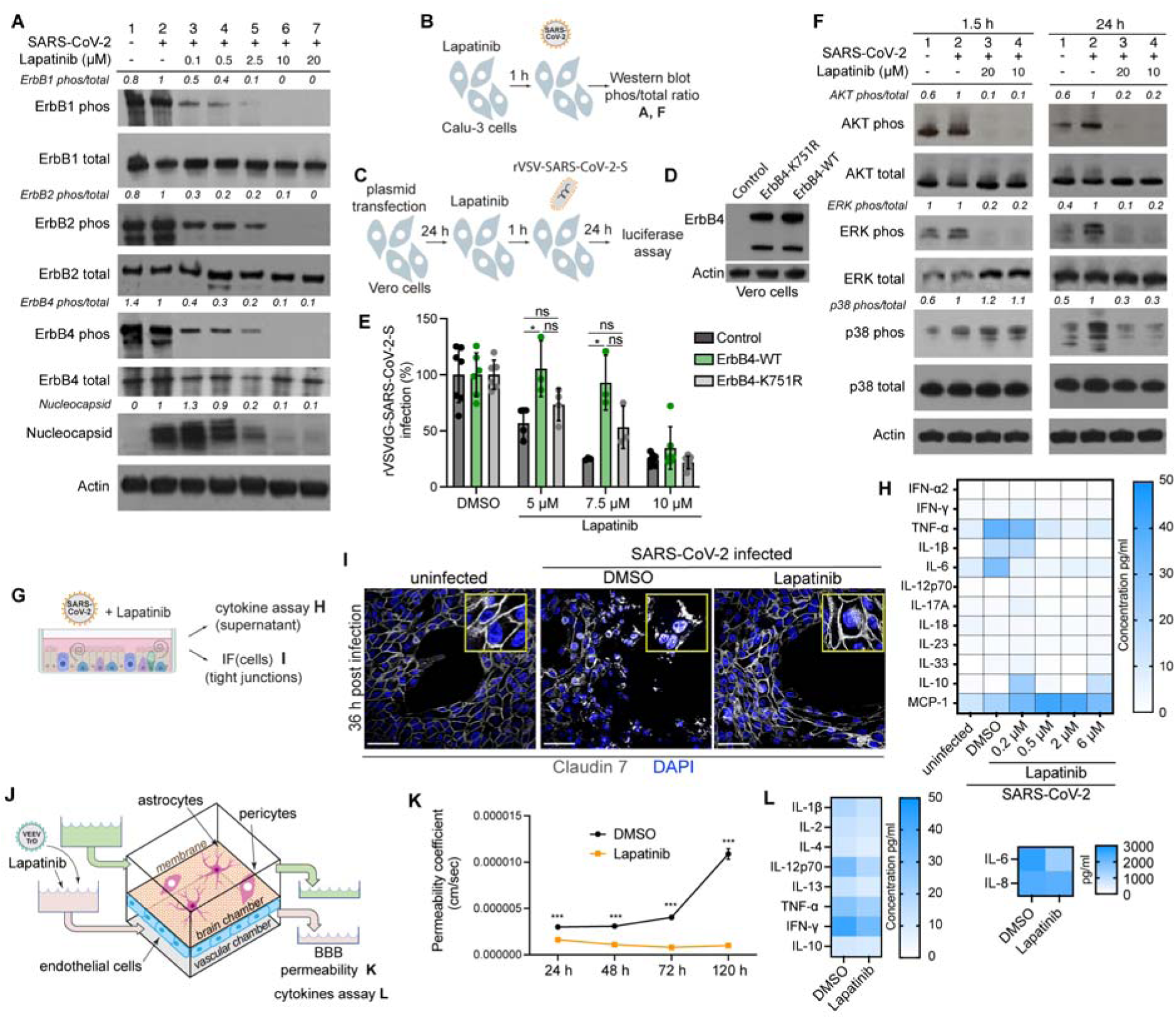
ErbBs are the molecular targets mediating the antiviral effect of lapatinib and they regulate viral-induced inflammation and tissue injury. **A, F** ErbBs (**A**), AKT, ERK, and p38 MAPK (**F**) phosphorylation and nucleocapsid expression (**A**) in Calu-3 cells that are uninfected (lane 1), infected and treated with DMSO (lane 2) or infected and treated with various concentrations of lapatinib (lanes 3-7) measured via Western blotting at 1.5 (**F**) and 24 (**A, F**) hpi with SARS-CoV-2 (USA-WA1/2020 strain, MOI=1). Shown are representative membranes blotted for phospho- and total proteins and quantitative phospho-to total protein ratio data relative to infected cells treated with DMSO (lane 2). **B, C** Schematics of experiments shown in A and F (**B**) and D and E (**C**). **D**, Level of ErbB4 and actin expression measured via Western blot following transfection of Vero cells with control or ErbB4-expressing plasmids. **E,** Rescue of rVSV-SARS-CoV-2-S infection in the presence of lapatinib upon ectopic expression of the indicated plasmids measured by luciferase assays 24 hpi in Vero cells. **G**, Schematic of the experiments shown in I and H. **H,** Heat map showing the concentration of cytokines (pg/mL) in ALO-derived monolayers’ supernatants under the indicated conditions at 48 hpi with SARS-CoV-2 measured by LEGENDplex (Biolegend) kit. **I,** Confocal IF microscopy images of Claudin 7 (grey) and DAPI (blue) in naïve or SARS-CoV-2-infected ALO-derived monolayers treated with DMSO or 10 µM lapatinib and imaged at 36 hpi. Representative merged images at 40x magnification are shown. Scale bars are 50 µm**. J**, Schematic of the experiments shown in K, L. **K**, Permeability of the endothelial layer of gNVUs infected with VEEV TrD and treated with lapatinib or DMSO assessed by quantification of FITC-dextran levels in longitudinal samples collected from the brain and vascular chambers. **L**, Heat map showing the concentration of cytokines (pg/mL) in the brain side of gNVUs treated with lapatinib or DMSO at 120 hpi with VEEV (TrD) measured by LEGENDplex (Biolegend) kit. Data in panels A, F, D, H are representative of two independent experiments. In panels E and K means±SD of results of two combined experiments (E) or one experiment (K) conducted each with three replicates are shown. \**P* < 0.05, \*\*\**P* < 0.001 relative to DMSO by one-way ANOVA followed by Tukey’s multiple comparisons test at each lapatinib concentration (E) or unpaired t test (K). Ns, non-significant.

To confirm that inhibition of ErbBs is a mechanism underlying the antiviral effect of lapatinib, we conducted gain-of-function “rescue” experiments in Vero cells infected with rVSV-SARS-CoV-2-S and U-87 MG cells infected with VEEV (TC-83). Ectopic expression of WT ErbB4, whose depletion suppressed both infections most prominently (**Supplemental Figure 5C, E**), either completely or partially reversed the antiviral effect of various concentrations of lapatinib on rVSV-SARS-CoV-2-S and VEEV (TC-83) infections (**Figure 6C-E, Supplemental Figures 7D-H**). In contrast, ectopic expression of a catalytically inactive ErbB4 mutant harboring a lysine to arginine substitution in position 751 (K751R) or control plasmid did not reverse the antiviral effect of lapatinib (**Figure 6E and Supplemental Figure 7F**). These findings validate ErbB4 as a mediator of the antiviral effect of lapatinib and indicate that its enzymatic activity is required for viral infection.

### ErbB inhibition suppresses viral-induced inflammation and tissue injury *ex vivo* in human adult lung organoid (ALO)-derived monolayers and gNVUs

Data from non-infectious ALI and ARDS animal and human models indicate that ErbB1 and 2 are key regulators of inflammation and tissue injury via activation of the p38 MAPK, AKT/mTOR and Ras/RAF/MEK/ERK pathways (22, 32–35). To test the hypothesis that these pathways are activated in SARS-CoV-2 infection and suppressed by the pan-ErbB inhibitory effect of lapatinib, we measured their activation in Calu-3 cells upon SARS-CoV-2 infection and/or lapatinib treatment by Western blot analysis. At 1.5 and 24 hpi, SARS-CoV-2 increased the ratio of phosphorylated to total protein level of AKT, ERK, and/or p38 MAPK by >1.5-2.5 fold (**Figure 6B, F**), in agreement with reports in other cell lines (36, 37). Lapatinib treatment dramatically inhibited SARS-CoV-2-induced activation of AKT and ERK both at 1.5 and 24 hpi and of p38 MAPK at 24 hpi (**Figure 6F**). In the more complex ALO-derived monolayer model, lapatinib treatment inhibited SARS-CoV-2-induced phosphorylation of AKT and ERK, albeit not p38 MAPK (**Supplemental Figure 7I**).

To further test the hypothesis that ErbB-regulated signaling mediates the inflammatory response to SARS-CoV-2 infection, we measured cytokine levels in ALO-derived monolayer supernatants upon SARS-CoV-2 infection and treatment with lapatinib or DMSO. SARS-CoV-2 infection increased the production of TNF-α, IL-1β and IL-6, in agreement with former reports (38). Lapatinib treatment dose-dependently reduced the expression level of these pro-inflammatory cytokines, with levels at or lower than those measured in uninfected ALO-derived monolayers achieved at drug concentration of 0.5 µM (**Figure 6G, H**). Concurrently, lapatinib increased the expression level of MCP-1, suggesting that it may augment innate immune responses (39).

To define the role of ErbB signaling in SARS-CoV-2-induced lung injury, we analyzed the effect of lapatinib on the integrity of tight junction formation in ALO-derived monolayers via confocal IF analysis. Claudin 7 staining of uninfected ALO-derived monolayers revealed a continuous membranous pattern (**Figure 6G, I and Supplemental Figure 7J**). Thirty-six hours following SARS-CoV-2 infection and DMSO treatment, claudin 7 stained as speckles or short segments that often appeared in the cytoplasmic region. This finding was accompanied by cell separation and destruction of the alveolar-like architecture. In contrast, ALO-derived monolayers treated with 10 µM of lapatinib exhibited intact claudin 7 morphology and subcellular distribution as well as preserved architecture of the alveolar-like structure, comparable to uninfected controls (**Figure 6G, I and Supplemental Figure 7J**).

To determine whether these observations are generalizable to other viral infections, we monitored the effect of lapatinib on BBB integrity in VEEV (TrD)-infected gNVUs by quantifying the permeability of FITC-dextran every 24 h for the total 120 h (**Figure 6J).** In the TrD-infected, DMSO-treated gNVUs, the permeability dramatically increased starting at 72 hpi. Contrastingly, the infected gNVUs treated with lapatinib (5 µM) maintained the integrity of the barrier (**Figure 6K**). In parallel, lapatinib treatment reduced the production of the proinflammatory cytokines IL-1β, IL-2, IL-4, IL-6, IL-8, IL-12p70, IL-13, and TNF-α, but not IFN-γ (antiviral) and IL-10 (immune suppressant), as measured via multiplexed ELISA in perfusion media from the brain compartment at 120 hpi (**Figure 6J, L)**.

Based on our and the cumulative published data, we propose a model wherein ErbBs are required for the life cycle of SARS-CoV-2 and VEEV, while pan-ErbB activation of downstream signaling pathways by these and other viruses mediates inflammation and tissue injury. By suppressing both processes, pan-ErbB inhibitors not only inhibit viral infection, but also protect from the resulting inflammation and the disruption of lung epithelium and BBB integrity (**Figure 7**). Since others have shown that lapatinib reverses increased epithelium permeability in a non-infectious model *in vitro* (32) and investigational ErbB inhibitors protect from acute and chronic lung injury in non-infectious models *in vivo* (33, 40–42), we propose that lapatinib’s protective effect from inflammation and tissue injury is only partly driven by its antiviral effect.

**Figure 7.**
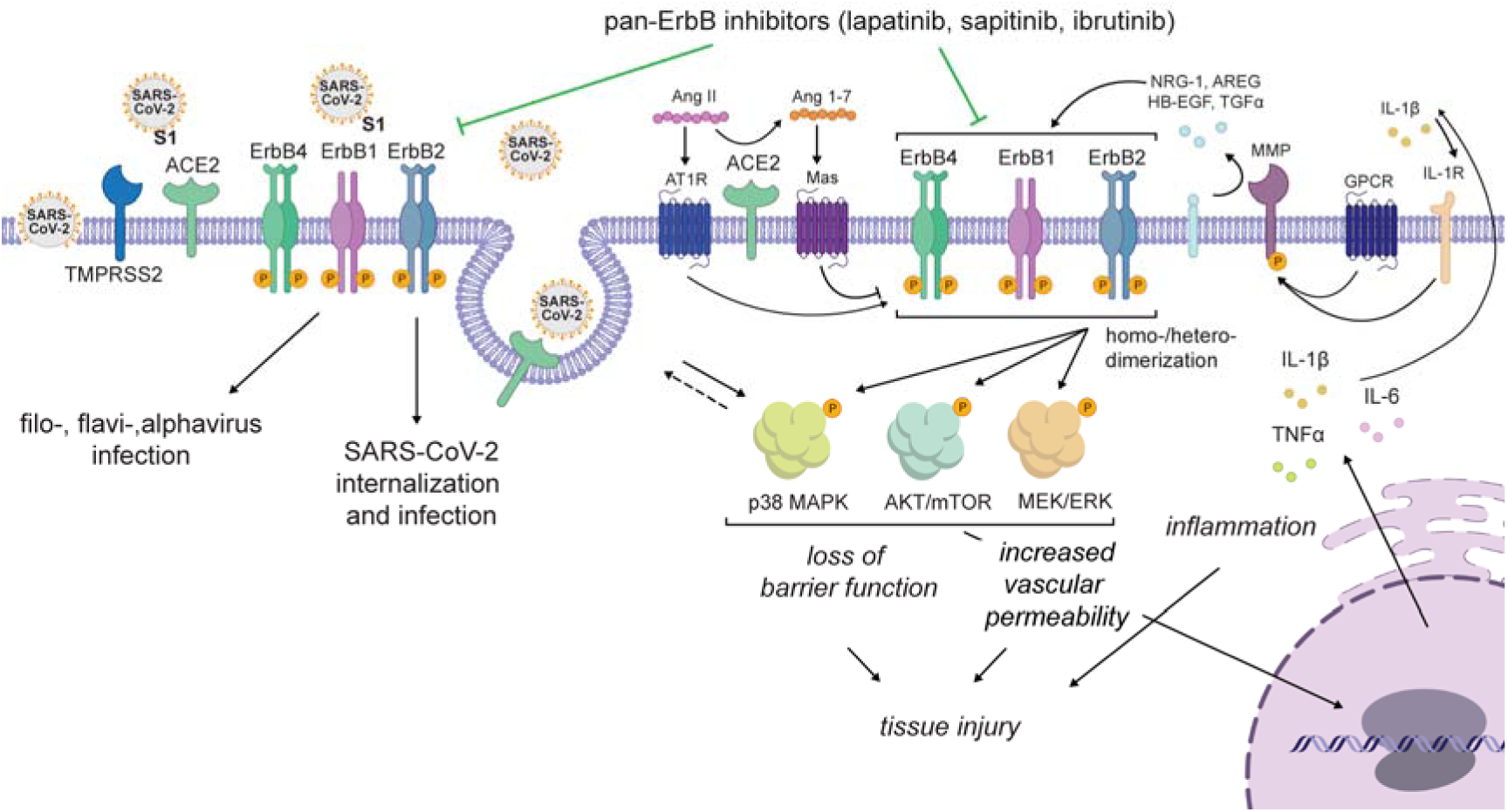
Proposed model for the roles of ErbBs in the regulation of viral infection and pathogenesis and the mechanism of action of pan-ErbB inhibitors. ErbBs regulate SARS-CoV-2 internalization and other stages of the viral life cycle and are required for effective replication of other emerging RNA viruses. Moreover, pan-ErbB activation promotes signaling in pathways implicated in inflammation and tissue injury in severe pandemic coronaviral infections and other disease model. By inhibiting ErbBs, lapatinib and other pan-ErbB inhibitors not only suppress viral infection but also protect from the resulting inflammation and tissue injury.

## Discussion

While ErbB1 has been implicated in the life cycle of multiple RNA and DNA viruses (43), its precise role in coronavirus infections, its role in alphavirus infections and the roles of ErB2 and ErbB4 in any viral infection remained unknown. Moreover, the relevance of ErbBs in viral-induced inflammation and acute tissue injury has not been reported. Here, we addressed this knowledge gap and studied the therapeutic potential of inhibiting ErbBs as a broad-spectrum antiviral strategy. Integrating virology, biochemical, genetic, immunological, and pharmacological approaches with *ex vivo* and *in vivo* models that recapitulate COVID-19 and VEEV pathology, we reveal regulation of SARS-CoV-2 and VEEV infections and subsequent inflammation and epithelial barrier or BBB injury by ErbBs, validating these kinases as attractive targets for antiviral therapy. Moreover, our findings provide a proof of concept for the utility of approved pan-ErbB inhibitors as broad-spectrum antiviral agents and reveal the mechanism of antiviral action and anti-inflammatory and tissue protective effects.

Using pan-ErbB inhibitors as pharmacological tools, we discover that ErbB1, ErbB2 and ErbB4 are required for effective SARS-CoV-2 and VEEV infections. Beyond pharmacologically, we genetically validate ErbBs as anti-SARS-CoV-2 and VEEV targets. Moreover, we provide multiple lines of evidence to support modulation of ErbBs activity as an important mechanism of the antiviral action of lapatinib. First, we show that lapatinib inhibits SARS-CoV-2 entry, analogous to the phenotype we reveal with RNAi-mediated suppression of ErbBs. Second, lapatinib’s antiviral activity correlates with reduced levels of phospho-ErbBs both *in vitro* and in ALO-derived monolayers. Third, WT, but not a kinase-dead ErbB4 mutant, reverses the anti-SARS-CoV-2 and anti-VEEV effect of lapatinib.

Our findings establish an effect of lapatinib on SARS-CoV-2 entry and provide insight into the roles of ErbBs in this stage of the viral life cycle. More specifically, lapatinib treatment reduced the colocalization of SARS-CoV-2 particles with late endosomes at early time points post-infection, revealing a role for ErbBs in viral internalization. Moreover, flow cytometry analysis revealed that lapatinib suppressed ACE2 internalization in uninfected cells and reversed the effect of SARS-CoV-2 infection on ACE2 but not NRP1 internalization. It is thus tempting to speculate that ErbBs regulate internalization of SARS-CoV-2 with ACE2. Our co-immunoprecipitation assays revealed interactions of the SARS-CoV-2 spike S1 subunit with ErbBs in the absence of ACE2 and NRP1, suggesting that beyond an ACE2-dependent mechanism, direct S1-ErbB binding may also play a role in viral entry. Indeed, there was a trend towards reduction of surface level expression of ErbB2 upon SARS-CoV-2 infection, suggesting that ErbB internalization may play a role in viral entry. Yet, the precise roles of these interactions and of ErbBs in viral entry represent important topics for further studies. Beyond entry, regulation of additional stages of the viral life cycle by ErbBs was suggested by the time-of-addition experiments. A recent HTS revealed that lapatinib inhibits the SARS-CoV-2 Mpro (44), proposing one potential mechanism for its post-entry effect.

We and others provide evidence that ErbBs mediate inflammation and lung injury. In non-infectious human and animal ALI/ARDS models, ErbBs are key regulators of inflammation, loss of epithelial barrier function, thrombosis, vasoconstriction, and the resulting fibrosis (22, 32–35)-- processes also involved in severe COVID-19 pathogenesis (11). Indeed, transcriptomic and phosphoproteomic studies revealed that activation of ErbBs and/or their downstream pathways are among the strongest detected upon infection of human cells with SARS-CoV-1 (45), SARS-CoV-2 (36, 37) and MERS (46), and in mice infected with SARS-CoV-1 (47). However, ErbB signaling has not been directly linked to coronavirus-induced inflammation and lung injury. We demonstrate SARS-CoV-2-induced activation of p38 MAPK, AKT, and ERK in human lung epithelium and ALO-derived monolayers and inhibition of phosphorylation of both ErbBs and these downstream effectors by lapatinib. Moreover, in human ALO-derived monolayers, we show that lapatinib treatment effectively suppresses SARS-CoV-2-induced secretion of pro-inflammatory cytokines and disruption of the lung epithelial barrier integrity. By inhibiting ErbB activation via multiple ligands implicated in lung injury--such as NRG-1, TGF-α, HB-EGF, and AREG, some of which have been shown to play a role in coronaviral infections (34, 48)-- lapatinib should, at least in theory, achieve a greater anti-inflammatory and tissue protective effect from approaches that target individual components of these pathways (e.g. antibodies targeting IL-1β, TGF-β and IL-6, and p38 MAPK inhibitors) (**Figure 7**). It is also intriguing to speculate that by restoring ACE2 levels on the surface of SARS-CoV-2-infected cells, lapatinib may help reverse the unopposed Angiotensin (Ang) II effect shown to activate ErbB pathways and increase pulmonary vascular permeability in animal models of non-viral lung injury (49, 50).

While it remains to be experimentally proven, since ErbB1 has been shown to be required for SARS-CoV-1 infection (51), and the pathways downstream of ErbBs are similarly upregulated in SARS-CoV-1- and MERS-infected cells, we predict that these findings may apply to other pandemic coronaviral infections. Lapatinib effectively suppressed the secretion of pro-inflammatory cytokines and loss of BBB integrity in the VEEV (TrD)-infected gNVU model, and protected mice from a lethal VEEV (TC-83) challenge, establishing that the proinflammatory and tissue protective effects of pan-ErbB inhibition are generalizable to other viral infections and tissues. Demonstrating these protective effects in the complex, biologically relevant, human ALO-derived monolayer and gNVU models as well as in a lethal mouse model elucidates the translatability of this approach. While lapatinib has not been studied for the treatment of viral infections to date, ibrutinib, a BTK inhibitor with potent pan-ErbB activity (23) that we show suppresses SARS-CoV-2 and VEEV infections, has shown protection from progression to severe COVID-19, albeit in a small number of patients (52).

There is an urgent need to establish a therapeutic portfolio for future pandemic preparedness. The design of broad-spectrum DAAs is challenged by the genetic diversity of viral species and replication strategies; however, a host-targeted approach could overcome this challenge. Lapatinib inhibits viruses from four unrelated viral families. Moreover, lapatinib demonstrates a higher barrier to resistance than a DAA, supporting the hypothesis that targeting host proteins that are not under the genetic control of viruses increases the barrier to resistance, in agreement with our findings with NUMB-associated kinase inhibitors (9). Simultaneous inhibition of several proviral kinases by a single drug (i.e. “polypharmacology”), in this case the three catalytically-active ErbBs and possibly STK10, RAF1, and RIPK2 whose depletion suppressed SARS-CoV-2 and/or VEEV infections, may further increase the effectiveness while minimizing viral resistance, as previously shown in cancer (53).

Remarkably, we provide evidence that lapatinib could achieve a synergistic effect with nirmatrelvir to inhibit SARS-CoV-2 infection. Beyond improving the antiviral effect, such synergy achieved by combining drugs with distinct mechanisms of action could enable dose reduction and may reduce the emergence of resistant mutations, as those already selected *in vitro* under paxlovid treatment and exist in clinical isolates (6, 7) underscoring the risk of broad administration of DAAs as monotherapies.

Repurposing existing drugs requires less capital and time and diminishes the clinical risks, as such drugs have already been tested (toxicity, pharmacokinetics (PK), dosing, etc.) for their primary indication (11). Lapatinib is an oral drug that is approved globally in combination drug treatments for metastatic, ErbB2-positive breast cancer. Based on the available PK data, the plasma level achieved with the approved dose of lapatinib (1500 mg once daily) in humans should be therapeutic as it is 8-10-fold higher than the EC_50_s we measured for its antiviral effect in ALO-derived monolayers and gNVUs. Even higher lapatinib lung levels may be achieved, as suggested by the predicted lung to plasma area under the curve ratio of 8.2-10 (54) and measured in mouse lungs (**Supplemental Figure 4E**). Although toxicity is a concern when targeting host functions, lapatinib has a favorable safety profile, particularly when used as a monotherapy and for short durations, as those required to treat acute infections. A summary of safety considerations and drug-drug interactions is provided in **Supplemental text 3**.

The other hits emerging from our HTS are discussed in **Supplemental text 4.**

In summary, our study validates ErbBs as druggable targets for antiviral, anti-inflammatory and tissue protective approaches and proposes approved drugs with anti-pan-ErbB activity as an attractive class of repurposing candidates for COVID-19 and VEEV that may provide readiness for future outbreaks of other emerging viruses.

## Online Methods

### Compounds, plasmids, and cells

Please see **Supplemental methods** for a complete list.

### HTS of compound libraries

Compounds were dispensed by an automated Agilent Bravo pipetting system in 384-well plates (Greiner #7810192) at a final concentration of 10 µM. As in a prior SARS-CoV-1 screening (55), Vero-E6-eGFP cells were plated in columns 1-24 24 hours before infection. 30 µL assay medium was added to columns 23 and 24 (cell controls). Following a 20-hour incubation, cells in columns 1-22 were infected with 30 µL SARS-CoV-2 (Belgium-GHB-03021) (MOI=0.001), using an automated, no-contact liquid handler (EVO100/Tecan) on the Caps-It robotics system. Plates were incubated for 4 days and imaged via a high content imager (Arrayscan XTI/Thermofisher). eGFP signal was used as a marker for survival. Cells were excited at 485-20 nm and emission was captured via a CCD camera and a BGRFRN_BGRFRN dichroic mirror and filter set. The exposure time was set at 0.023 seconds. Imaging acquisition speed was optimized using a 2×2 binning on 1104×1104 pixel resolution and reducing the number of autofocus focal planes. The Cellomics (Thermofisher) software was used for image analysis. A custom-made image analysis protocol was created using the SpotDetector bioapplication. SpotTotalAreaCh2 was used for further data analysis.

### Viral stocks preparation and sequencing

Belgium-GHB-03021 SARS-CoV-2 strain was recovered from a nasopharyngeal swab taken from a patient returning from China early February 2020 (56) and passaged 6 times on Huh7 and Vero E6 cells. 2019-nCoV/USA-WA1/2020 SARS-CoV-2 isolate (NR-52281) (BEI Resources) was passaged 3-6 times in Vero E6-TMPRSS2 cells. The rSARS-CoV-2/WT and rSARS-CoV-2/Nluc (rSARS-CoV-2 expressing Nluc-reporter gene) viral stocks were generated as previously described (ref) (57). USA-WA1/2020 from passage 3 used for the majority of the experiments was subject to SARS-CoV-2 whole-genome amplicon-based sequencing on a MiSeq platform (Illumina) by adapting an existing pipeline as described in (58), showing no deletion or point mutations in the multi-basic cleavage (MBC) domain. Belgium/GHB-03021/2020 SARS-CoV-2 from passage 6 was sequenced following a metagenomics pipeline (59) showing 100% deletion of the MBC domain. VEEV-TC-83-nLuc RNA was transcribed *in vitro* from cDNA plasmid templates linearized with MluI via MEGAscript SP6 kit (Invitrogen #AM1330) and electroporated into BHK-21 cells. DENV RNA was transcribed *in vitro* from pACYC-DENV2-NGC plasmid by mMessage/mMachine (Ambion) kits and electroporated into BHK-21 cells. WT Trinidad Donkey (TrD) VEEV strain, EBOV (Kikwit isolate) and MARV (Ci67 strain) (BEI Resources) were grown in Vero E6 cells. Supernatants were collected, clarified and stored at -80 °C. Viral titers were determined via standard plaque assays on BHK-21 (DENV, VEEV) or Vero E6 cells (SARS-CoV-2, EBOV, MARV).

### rVSV-SARS-CoV-2-S production

HEK-293T cells were transfected with spike expression plasmid followed by infection with VSV-G pseudotyped ΔG-luciferase VSV virus and harvesting of culture supernatant, as described (58).

### Human adult lung organoids (ALOs)

The ALO model was generated from adult stem cells isolated from deep lung biopsy specimens (13). This model is complete with all 6 cell types of proximal and distal airways as validated previously (13). Lung-organoid-derived monolayers were prepared (13) and plated in Pneumacult Ex-Plus Medium (StemCell, Canada).

### gNVUs model

NVUs were prepared as described in (17, 18)

### Infection assays and pharmacological inhibition

Unless stated otherwise, inhibitors or DMSO were added to the cells 1 hour prior to viral inoculation and were left for the duration of the experiment. Calu-3 cells, Vero cells or ALOs were infected with SARS-CoV-2 in triplicates (MOI=0.05 or 1) in DMEM containing 2% FCS at 37°C under biosafety level 3 (BSL3) conditions. After 1 to 3-hour incubation, the inoculum was removed, cells were washed and supplemented with new medium. Culture supernatants were harvested for measurement of viral titer by standard plaque assays and cells were lysed in TrizolLS for RT-qPCR analysis. Huh7 cells were infected with DENV2 in replicates (n=3-10) (MOI= 0.05). At 48 hpi infection was measured via luciferase or plaque assays. Huh7 cells were infected with EBOV (MOI=1) or MARV (MOI=2) under BSL4 conditions. At 48 hpi, cells were formalin-fixed for 24 hours prior to removal from BSL4. Infected cells were detected using an EBOV or MARV glycoprotein-specific monoclonal antibody (KZ52 and 7E6, respectively) and quantitated by automated fluorescence microscopy using an Operetta High Content Imaging System (PerkinElmer). U-87 MG cells were infected with VEEV-TC-83-nLuc in 5 replicates (MOI=0.01) or with WT VEEV (TrD) in triplicates. At 24 hpi, infection was measured via nanoluciferase or plaque assays. gNVUs were perfused with lapatinib- or DMSO-containing culture medium for 1 hour followed by introduction of medium containing VEEV-TrD (MOI=0.1) into the vascular inlet and 1-hour incubation at 37°C. Lapatinib-containing medium was reintroduced into the gNVU daily and the units were maintained at 37°C, 5% CO_2_ conditions for up to 120 hours (study duration).

### Viability assays

Viability was assessed using alamarBlue reagent (Invitrogen) according to the manufacturer’s protocol. Fluorescence was detected at 560 nm on an InfiniteM1000 plate reader (Tecan).

### Combination drug treatment

Calu-3 cells were treated with lapatinib-DAAs combinations and infected with rSARS-CoV-2/Nluc (USA-WA1/2020 strain) (MOI=0.05). Twenty-four hpi the antiviral effect was measured via Nluc assay and cellular viability was measured via alamarBlue assay. The MacSynergy II program was used for data analysis, as described (9, 14, 60). Matrix data sets in three replicates were assessed at the 95% confidence level for each experiment.

### RNA interference

siRNAs (10 pmol/96-well) were transfected into cells using lipofectamine RNAiMAX transfection reagent (Invitrogen) 48 hours prior to viral infection. ON-TARGETPlus siRNA SMARTpools against 7 genes and non-targeting siRNA (siNT) were purchased from Dharmacon/Horizon Discovery with gene IDs as follows: EGFR (1956), ErbB2 (2064), ErbB4 (2066), RIPK2 (8767), RAF1 (5894), STK10 (6793), MAP2K5 (5607).

### Time-of-addition assay

Calu-3 cells were infected with SARS-CoV-2 (MOI=1). Two hpi, virus was removed, and cells were washed twice with PBS. At distinct time points, 10 µM lapatinib or 0.1% DMSO were added. Cell culture supernatants were collected at 10 hpi, and infectious viral titers were measured by plaque assay.

### Temperature-shift assay

Vero cells were either concurrently inoculated with rVSV-SARS-CoV-2-S and treated with lapatinib (10 µM) or DMSO at 4°C for 2 hours and washed, or infected with the virus at 4°C for 2 hours, washed, and then treated with lapatinib (10 µM) or DMSO at 37°C for 2 hours. Entry was measured at 24 hpi.

### Entry assays

At 48 h after siRNA transfection, Calu-3 cells were infected with WT SARS-CoV-2 (MOI=1). At 1 hpi, cells were washed three times with PBS and fresh medium added. At 2 hpi, cells were lysed in TRIzolLS (Invitrogen) and intracellular viral RNA levels were measured by RT-qPCR.

### RT-qPCR

RNA was extracted from cell lysates using Direct-zol RNA Miniprep Plus Kit (Zymo Research) and reverse transcribed using High-Capacity cDNA RT kit (Applied Biosystems). Primers and PowerUp SYBR Green Master Mix (Applied Biosystems) were added to the samples, and PCR reactions were performed with QuantStudio3 (Applied Biosystems) in triplicates. Target genes were normalized to GAPDH. Sequences of primers used for RT-qPCR are available upon request.

### Gain-of-function assays

Plasmids encoding ErbB4 or control were transfected into cells using Lipofectamine 3000 reagent (Invitrogen) 24 hours prior to drug treatment and viral infection. Viral infection and cell viability were measured 24 hours later via luciferase and alamarBlue assays, respectively.

### Resistance studies

VEEV (TC-83) was used to inoculate U-87 MG cells (MOI=0.1) and passaged daily under increasing drug selection (2.5-5 μM, passages 1–3; 5-10 μM, passages 4-7; 10-15 μM, passages 8-10). Following 10 passages, viral titers were measured in culture supernatants by plaque assays. ML336 resistant mutation emerging in nsP2 at passage 10 was confirmed by purification and reverse transcription of viral RNA from cell supernatants using RNeasy Mini Kit (Qiagen) and SuperScript IV First-Strand Synthesis kit (Invitrogen). The nsP2 region was amplified with Platinum Green Hot Start PCR Master Mix (2x) (Invitrogen) using the following primers: (forward: AGGAAAATGTTAGAGGAGCACAAG reverse: GTCAATATACAGGGTCTCTACGGGGTGT) and sequenced (Sequetech Corp.).

### *In vitro* kinase assays

These assays were performed on the LabChip platform (Nanosyn) or radiometric HotSpot^TM^ platform (Reaction Biology).

### Signaling pathway analysis

Following 2-hour starvation, Calu-3 cells or ALO-derived monolayers were treated with lapatinib or DMSO and within an hour infected with SARS-CoV-2 (MOI=1). Cell lysates were obtained at 1.5 and/or 24 hpi followed by Western blot analysis with antibodies targeting the phosphorylated and total protein forms: anti-ErbB4 (Santa Cruz), ErbB2 (Cell Signaling), ErbB1, AKT, ERK, p38 (Cell Signaling), P-ErbB4 (Tyr1284), P-ErbB2 (Tyr1248), P-ErbB1 (Tyr1173), P-AKT (Ser473), P-ERK (Thr202/Tyr204), P-p38 (Thr180/Tyr182) (Cell Signaling), and β-actin (Sigma-Aldrich, catalog A3854) antibodies. Phosphorylated to total protein ratios were quantified with ImageJ software (NIH).

### Cytokine measurements

Cytokine concentrations in ALO supernatants were quantified using LEGENDplex Human Inflammation Panel 1 (Biolegend) kit and read on a Quanteon (Agilent). Data was analyzed using LEGENDplex V8.0 software. Cytokine levels in gNVU perfused media were quantified using the MSD V-PLEX Proinflammatory Panel Human Kit (Cat. K15049D-2) and MESO QuickPlex SQ 120 (MesoScale Discovery) reader.

### BBB Permeability Assay

A 3kD FITC dextran assay (ThermoFisher Scientific) was used and the effective permeability of the brain microvascular endothelial cells monolayer was calculated, as described (18).

### Animal Studies

C3H/HeN female mice (6-8 weeks of age) (n=5 per group) were inoculated intranasally with 5×10^6^ PFU of VEEV (TC-83). Animals were pretreated for 12 hours and post treated once or twice a day with lapatinib (200 mg/kg) in 0.5% hydroxypropyl methylcellulose with 0.1% Tween-80 as vehicle through oral gavage. Survival was monitored up to a period of 2 weeks. Control mice were TC-83 infected and treated with vehicle only. In each group, three animals were sacrificed at days 3, 7 and 10 post-infection and virus titers were measured in both serum and brain by plaque assays. Mice were euthanized with CO_2_ and the serum was collected with 23-gauge needle right away, by cardiac puncture, 500–700 ml of blood was collected from each mouse, the blood was spun for 5 minutes at 14000 rpm, serum was collected, and viral plaque analysis was performed. Mouse brains were collected from euthanized animals, homogenized in DMEM supplemented media using IKA ULTRA-TURRAX Tube drive and DT-20 tube system and used for plaque assays. Multi-dose PK study in C57BL/6 mice was conducted by Sai Life Sciences (India).

### Immunofluorescence and confocal microscopy

ALO-derived monolayers were washed with PBS, fixed, blocked, and incubated with mouse mAb SARS-CoV-2 nucleocapsid antibody (SinoBiological) and rabbit Claudin 7 polyclonal antibody (ThermoFisher) overnight at 4°C, followed by incubation with secondary antibodies, and counterstaining with DAPI (ThermoFisher) and phalloidin (ThermoFisher). Images were taken on an SP8 microscope (Leica).

For imaging viral particles, VeroE6-TMPRSS2 cells were pretreated with lapatinib (10 µM) or DMSO for 1 hour at 37°C and infected with SARS-CoV-2 (MOI=1) at 4°C. Following 1-hour incubation, infected cells treated with lapatinib or DMSO were subject to temperature shift to 37°C to initiate virus infection. At 1 hpi, cells were washed and fixed as described above. Cells were incubated with mouse mAb SARS-CoV-2 nucleocapsid antibody (SinoBiological) and Rab7 (Origene, AB0033-200) followed by a secondary antibody. Images were taken using LSM 880 microscope (Zeiss) with 63x objective. Colocalization was quantified using ImageJ (JACoP) colocalization software and Manders’ colocalization coefficients (MCCs).

### Co-immunoprecipitations

A549-NRP1^KO^ cells were co-transfected with plasmids expressing S1-Flag and ErbB1, ErbB2 or ErbB4 using Lipofectamine 3000 reagent. At 24 h post-transfection, the cells were lysed with M-Per protein extraction reagent (Thermo Fisher Scientific). Clarified supernatants were precleared with Dynabeads protein G (Invitrogen) for 1 hour at 4°C and incubated with either anti-ErbB1, ErbB2, ErbB4 (Cell Signaling) or IgG antibody overnight at 4°C. The antibodies and bound proteins were captured by protein G Dynabeads for 2 hours at 4°C. Beads were washed and resuspended in SDS sample buffer.

### Flow cytometry

Calu-3 cells were pretreated with lapatinib (10 µM) or DMSO for 1 hour at 37°C and infected with SARS-CoV-2 (MOI=1) at 4°C. Following 1-hour incubation, the temperature was shifted to 37°C to initiate infection. At 0.5 and 1 hpi, cells were washed with PBS and incubated for 15 min at RT with Zombie Aqua™ live/dead fixable dye (Biolegend) and FcR Blocking Reagent (Miltenyi Biotec). Cells were stained for 20 min at 4°C with the following antibodies: rat anti-human NRP1-BV421 (Biolegend), goat anti-human ACE2-APC (R&D systems) and mouse-anti human ErbB2-Alexa Fluor 488 (R&D systems), or with their corresponding isotype controls. Unbound antibody was washed, and cells were fixed with 4% PFA for 1 hour at RT. Cell acquisition was performed on an Aurora Cytek spectral flow cytometer and data was analyzed using FlowJo software (TreeStar).

### Statistics

Data were analyzed with GraphPad Prism software. EC_50_ and CC_50_ values were measured by fitting of data to a 3-parameter logistic curve. *P* values were calculated by one-way ANOVA with either Dunnett’s or Tukey’s multiple comparisons tests or by Student’s t-test.

### Study Approval

SARS-CoV-2, VEEV and filovirus work were conducted in BSL3 and BSL4 facilities at Stanford University, KU Leuven Rega Institute, GMU, and USAMRIID according to CDC and institutional guidelines. Human lung organoids propagation was approved under protocol IRB# 190105 at UCSD. Animal experiments were approved by the Institute of Laboratory Animal Resources, National Research Council, NIH Publication No. 86–23.

## Acknowledgements

This work was supported by awards number W81XWH2210283 and W81XWH-16-1-0691 from the Department of Defense (DoD)/Congressionally Directed Medical Research Programs (CDMRP) (to SE and SDJ), award number HDTRA11810039 from the Defense Threat Reduction Agency (DTRA)/Fundamental Research to Counter Weapons of Mass Destruction (to SE, SDJ and AN), grants number 1 R01 AI 158569–01 (to SE and SDJ), 3R01DK107585-05S1 (to SD), UCOP-R00RG2642 (to SD and PG), and UCOP-R01RG3780 (to PG and DS), and 3UH3TR002097-04S1 (to JPW and JAB) from the National Institutes for Health (NIH), and a gift from The Frank and Denise Quattrone Foundation (to SE). S.E. is a Chan Zuckerberg Biohub investigator. M.K. was supported by a Postdoctoral Fellowship in Translational Medicine by the PhRMA Foundation. L.G. was supported by a long-term European Molecular Biology Organization (EMBO) Fellowship (ALTF 584-2021). V. D was supported by a Chan Zuckerberg Biohub Collaborative Postdoctoral Fellowship.

Flow cytometry analysis and IF confocal microscopy were done on instruments in the Stanford Shared FACS Facility and Stanford University Cell Sciences Imaging Core Facility (RRID:SCR_017787) respectively. We thank the Stanford SPARK program’s advisors for critical advice and investigators who have provided plasmids (see Methods). We thank the Stanford *in vitro* BSL3 service Center and its Director Dr. Jaishee Garhyan and Dr. Arjun Rustagi for their assistance in the BSL3. We also thank the staff of Stanford Clinical Virology Laboratory for their help sequencing the SARS-CoV-2 USA-WA1/2020 isolate used in this work. We thank Clayton M. Britt and David K. Schaffer at the Vanderbilt Microfabricated Technologies Resource for the care with which they fabricated the gNVUs. Opinions, conclusions, interpretations, and recommendations are those of the authors and are not necessarily endorsed by the funders. The mention of trade names or commercial products does not constitute endorsement or recommendation for use by the Department of the Army or the Department of Defense.

## Author Contributions

S.S., M.K., L.G., P-T.H., W.C., V.D., C-W.L., S.K., N.B., P.L., F. A., N.A.B., D.H.N.T., C.A.C, K.E.H., M.S. designed and performed the experiments and conducted data analysis. J.A.B., C.T., C.Y., A.M.K., P.G. S.D., D.E.S-C., J.J., J.P.W., provided reagents and guidance. S.E., S.D.J., D.J., D.E.S-C., J.N., A.N, B.A.P, and L.M-S. provided scientific oversight and guidance. S.S., S.E., S.D.J., M.K., W.C., J.M.D., and D.J. wrote the first version of the manuscript. S.E., D.J., J.N., A.N., S.D.J., J.M.D. provided funding for the studies.

## Supplemental data

**This document includes:**

1. Supplemental figures and figure legends

2. Supplemental texts

3. Supplemental methods

4. Supplemental data references

### Supplemental figures

**Supplemental Figure 1:**
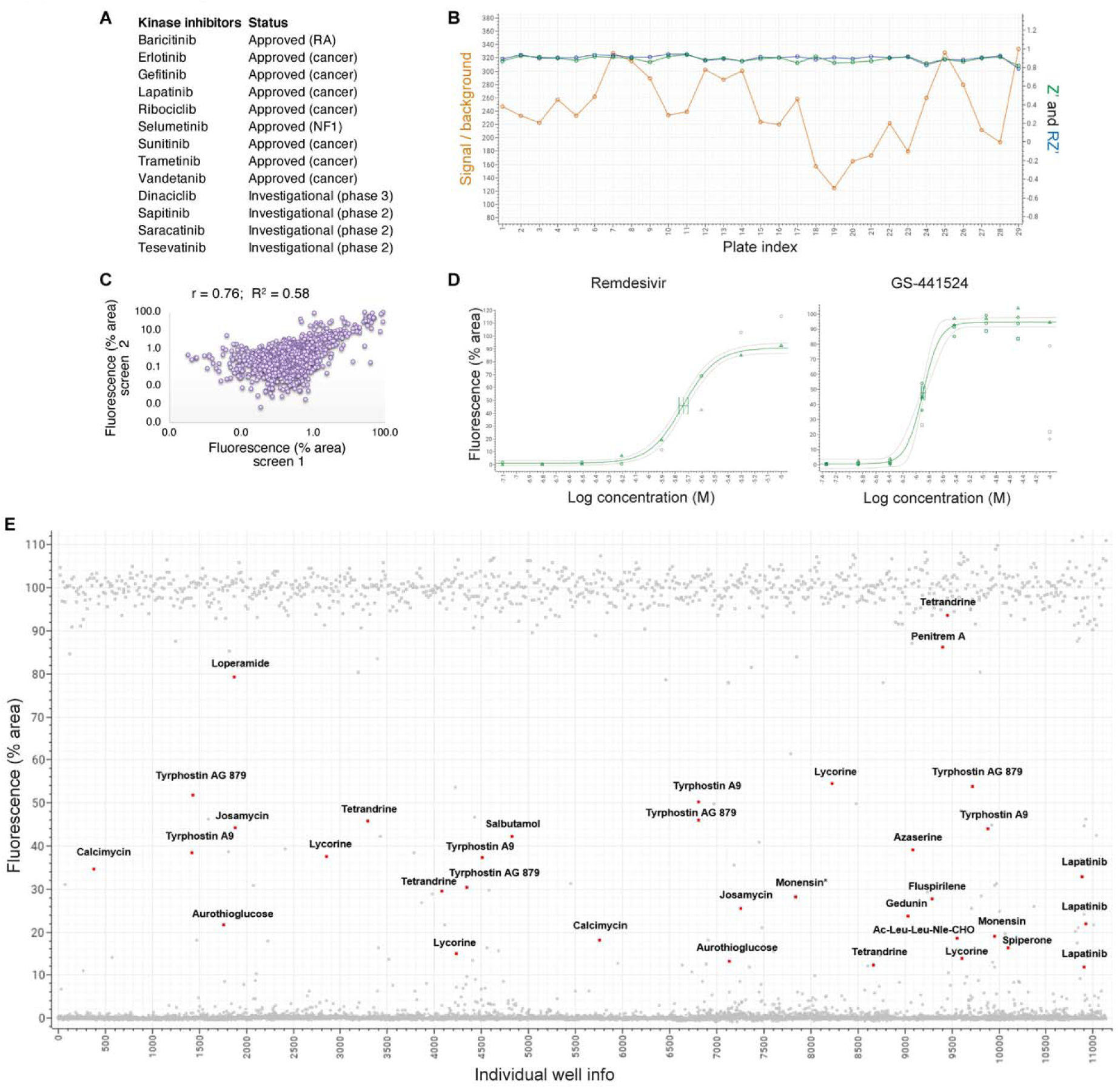
Characteristics and of the HTS. Related to figure 1. **A,** The kinase inhibitors included in the self-assembled set. **B,** Quality control of each individual plate of the 29 screened by determination of the signal-to-background (S/B), and the Z’ and RZ’ values. All three parameters were measured for each 384-well screening plate using the virus control (infected, DMSO treated) and cell control (uninfected, untreated) wells. S/B values ranged from 124 – 333. Z’ and RZ’ values were > 0.78. Generally, S/B values >10 and (R)Z’ values >0,5 are accepted as qualitative assays. All parameters were calculated using Genedata Screener. **C,** Scatter plot of the two replicate screens with a Pearson’s correlation coefficient (*r*) of 0.76 and R^2^ 0.58. **D**, Dose-dependent rescue of Vero-eGFP cells from SARS-CoV-2-induced lethality by remdesivir and its major active metabolite, GS-441524, used as positive controls, 4 days post-infection with SARS-CoV-2 (Belgium-GHB-03021, MOI=0.001). **E**, Percentage of fluorescence area values from all wells including the virus controls (infected, DMSO treated) and the cell controls (uninfected, untreated) from the 29 384-well plates. The red dots depict hits emerging in the screening. Grey dots represent reference compounds such as nelfinavir, GS-441524 and compounds not prioritized for further analysis.

**Supplemental Figure 2.**
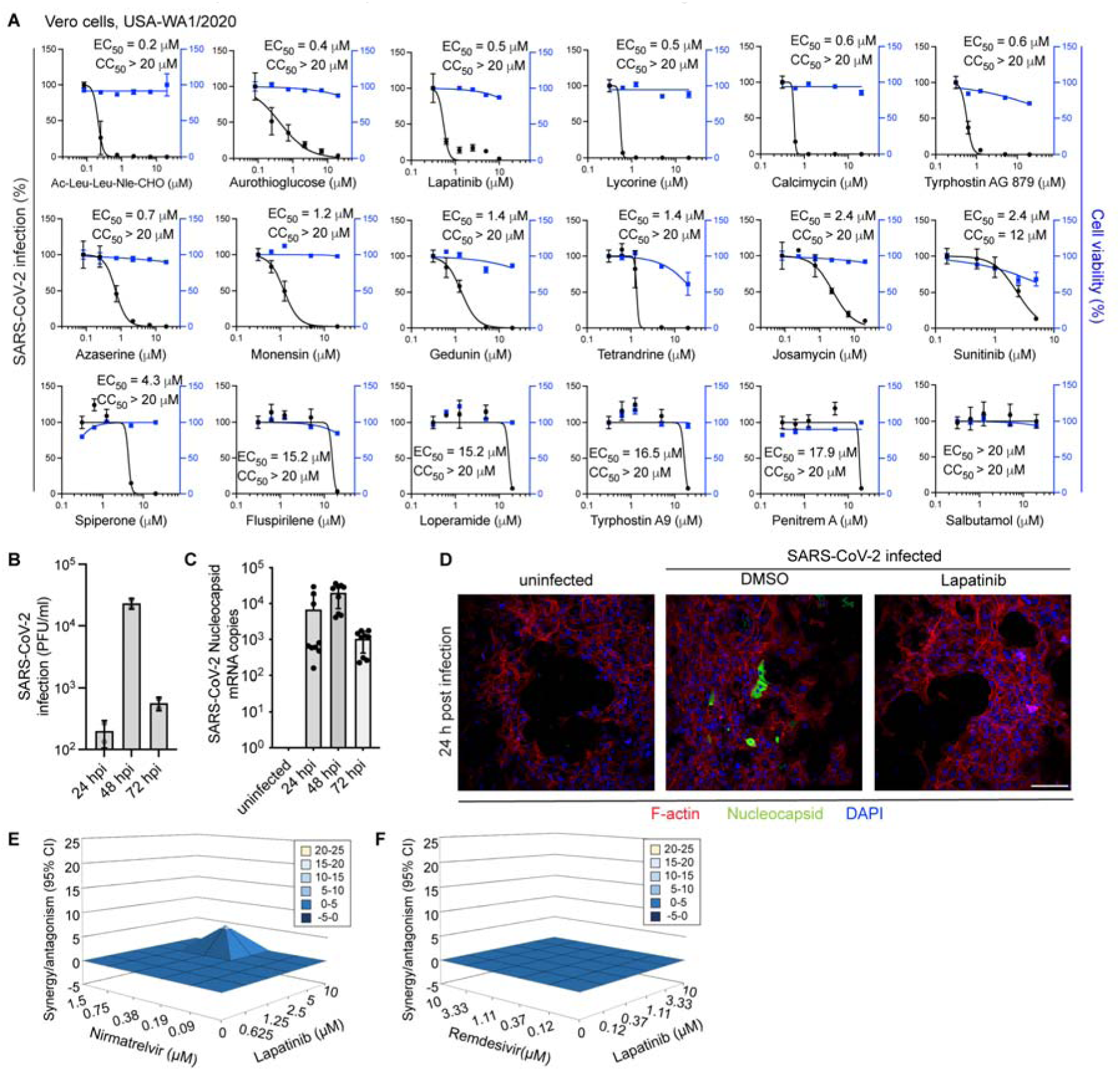
Validation of hits emerging from the HTS and characterization of human ALO-derived monolayers for studying the antiviral effect of emerging hits. Related to figures 1 and 2. **A,** Dose response curves to the indicated hits emerging from the HTS of SARS-CoV-2 infection (black, USA-WA1/2020 strain, MOI=0.05) and cell viability (blue) in Vero cells measured via plaque and alamarBlue assays at 24 hpi, respectively. **B, C** Viral titer by plaque assays in culture supernatants (**B**) and viral nucleocapsid (N) copy number analyzed by RT-qPCR in lysates (**C**) from human ALO-derived monolayers at 24, 48 and 72 hpi. **D**, Confocal IF microscopy images of F-actin (red), SARS-CoV-2 nucleocapsid (green) and DAPI (blue) in naïve and SARS-CoV-2-infected ALO-derived monolayers pre-treated with DMSO or 10 µM lapatinib at 24 hpi. Representative merged images at 20x magnification are shown. Scale bar is 100 μm. **E, F**, Synergy/antagonism of lapatinib and nirmatrelvir (**E**) or remdesivir (**F**) combination treatment on cellular viability measured in Calu-3 cells infected with rSARS-CoV-2/Nluc (USA-WA1/2020 strain) at 24 hpi via alamarBlue assays. Data represent differential surface analysis at the 95% confidence interval (CI), analyzed via the MacSynergy II program. Synergy and antagonism are indicated by the peaks above and below the theoretical additive plane, respectively. The level of synergy or antagonism is depicted by the color code. Data are representative of two independent experiments with three replicates each.

**Supplemental Figure 3.**
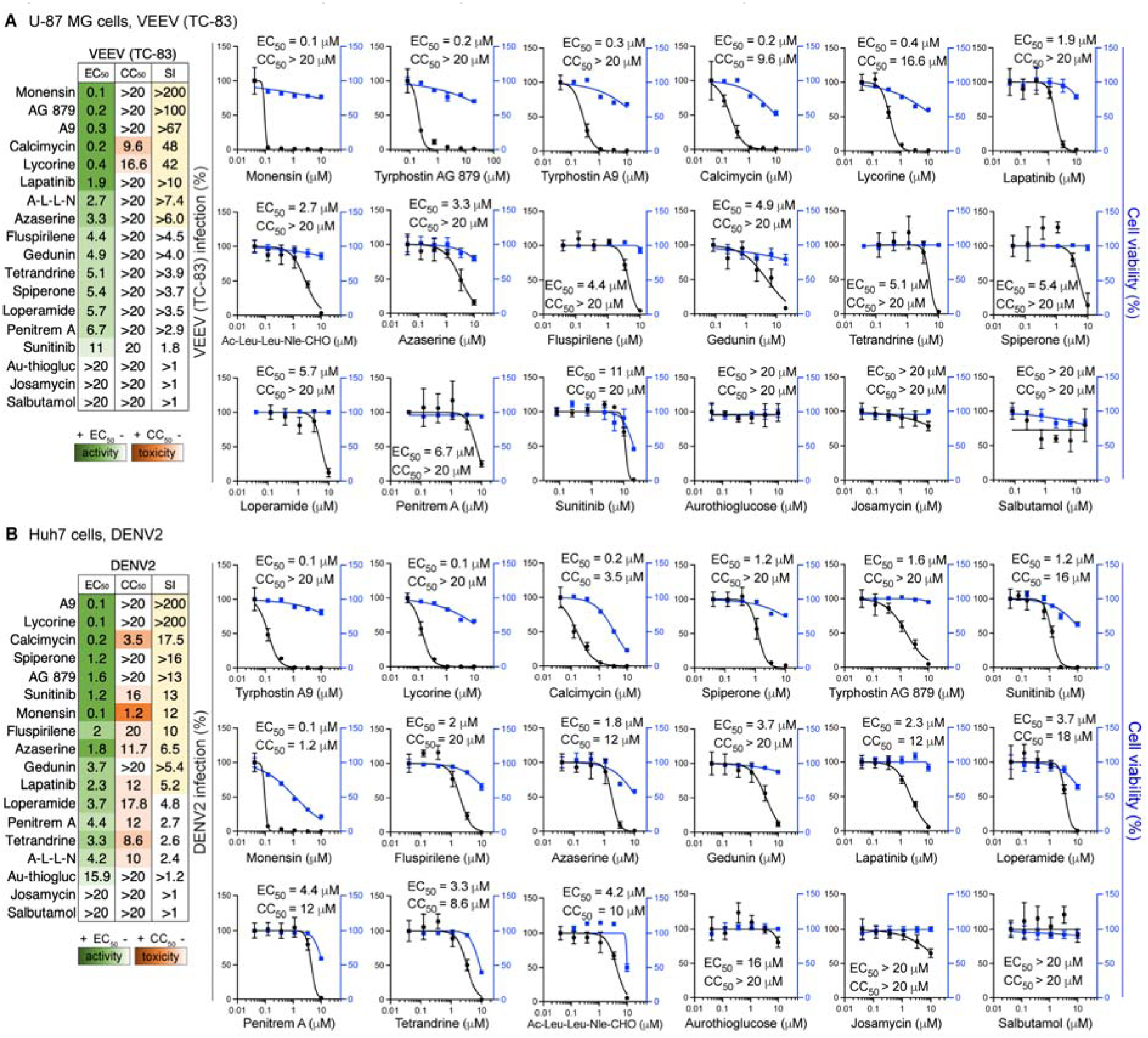
Broad-spectrum potential of hits. Related to figure 1,2. **A, B,** The 18 compounds emerging from the HTS were tested for their effect on VEEV (TC-83) (**A**) and DENV2 (**B**) infections in U-87 MG and Huh7 cells, respectively, via luciferase assays, and for their effect on cell viability via alamarBlue assays. Left panels: Heat maps of the EC_50_ and CC_50_ values of the indicated compounds color-coded based on the antiviral activity (green) and toxicity (orange). Selectivity indices (SI) greater than 5 are depicted in yellow. Right panels: Dose response curves to the indicated compounds of VEEV (TC-83) (MOI=0.1) or DENV2 (MOI=0.05) infections (black) in U-87 MG and Huh7 cells, respectively, measured via luciferase assays and cell viability (blue) measured by alamarBlue assays at 24 hours post-infection.

**Supplemental Figure 4:**
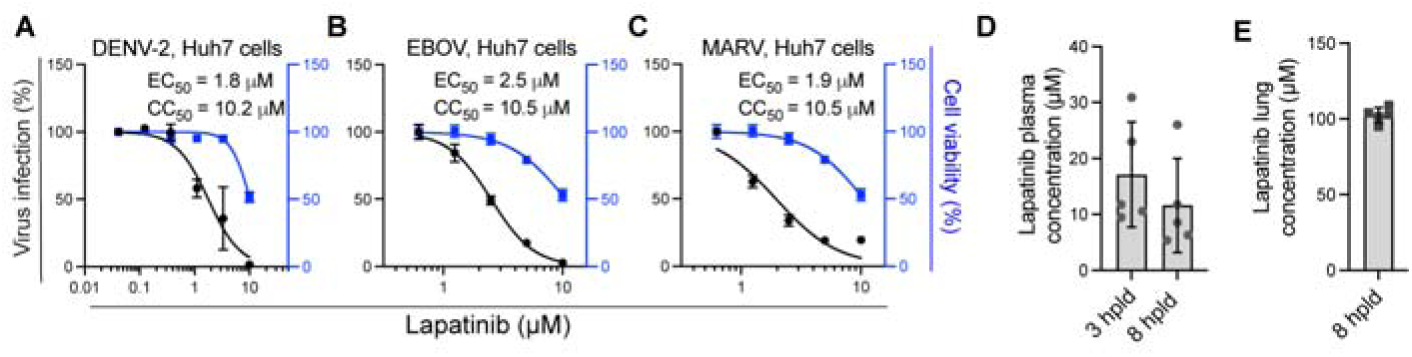
Lapatinib is a potent broad-spectrum antiviral. Related to figure 3. **A,** Dose response of DENV2 infection (blue) and cellular viability (black) to lapatinib measured in Huh7 cells via plaque and alamarBlue assays at 24 hpi (MOI=0.1), respectively. **B, C** Dose response of EBOV (Kikwit isolate, MOI=1) (H) and MARV (Ci67 strain, MOI=2) (I) infections (blue) and cellular viability (black) to lapatinib measured in Huh7 cells 48 hpi via microneutralization assay and CellTiter-Glo luminescent cell viability assay, respectively. **D, E** lapatinib’s plasma (**D**) and lung (**E**) concentrations after 8 days of twice daily treatment with 200 mg/kg in C57BL/6 mice measured 3 (E) and 8 (E, F) hours post last dose (hpld).

**Supplemental Figure 5:**
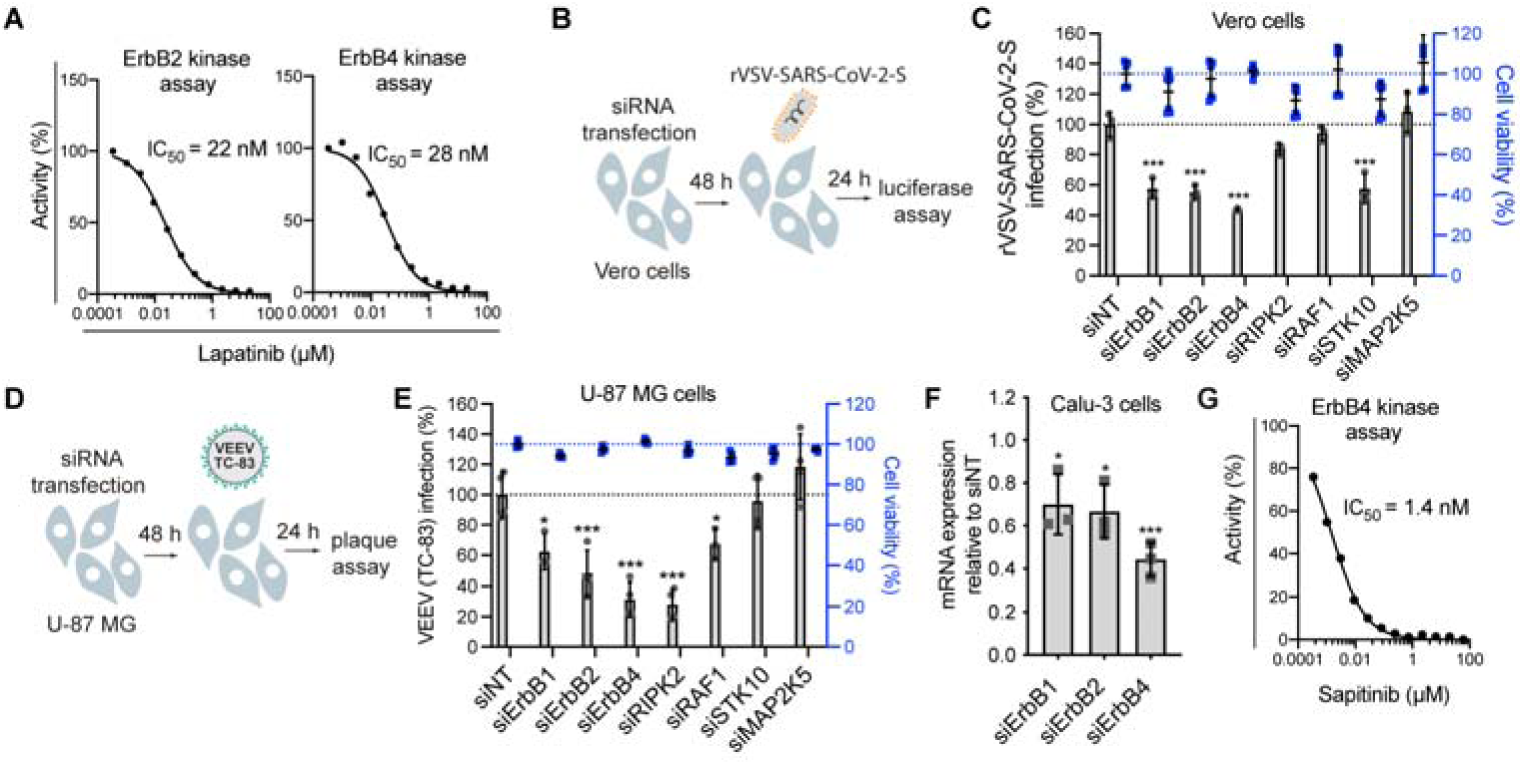
Validation of ErbBs as an antiviral target. Related to figure 4. **A**, Dose response to lapatinib of ErbB2 and/or ErbB4 kinase activity *in vitro* (Nanosyn). **B**, Schematic of the experiment shown in panel C. **C,** Percentage of infection by luciferase assays (grey) and cell viability by alamarBlue assays (blue) measured at 24 hpi of Vero cells transfected with the indicated siRNA pools with rVSV-SARS-CoV-2-S pseudovirus. **D**, Schematic of the experiment shown in panel E. **E**, Percentage of infection by plaque assays (grey) and cell viability by alamarBlue assays (blue) measured at 24 hpi of U-87 MG cells transfected with the indicated siRNA pools with VEEV (TC-83). **F**, Confirmation of siRNA-mediated gene expression knockdown by RT-qPCR in Calu-3 cells. Shown is gene expression normalized to GAPDH and expressed relative to the respective gene level in the siNT control at 48 hours post-transfection. **G,** Dose response to sapitinib of ErbB4 kinase activity *in vitro* (Nanosyn). \**P* < 0.05, \*\*\**P* < 0.001 relative to siNT by one-way ANOVA followed by Dunnett’s multiple comparisons test.

**Supplemental Figure 6:**
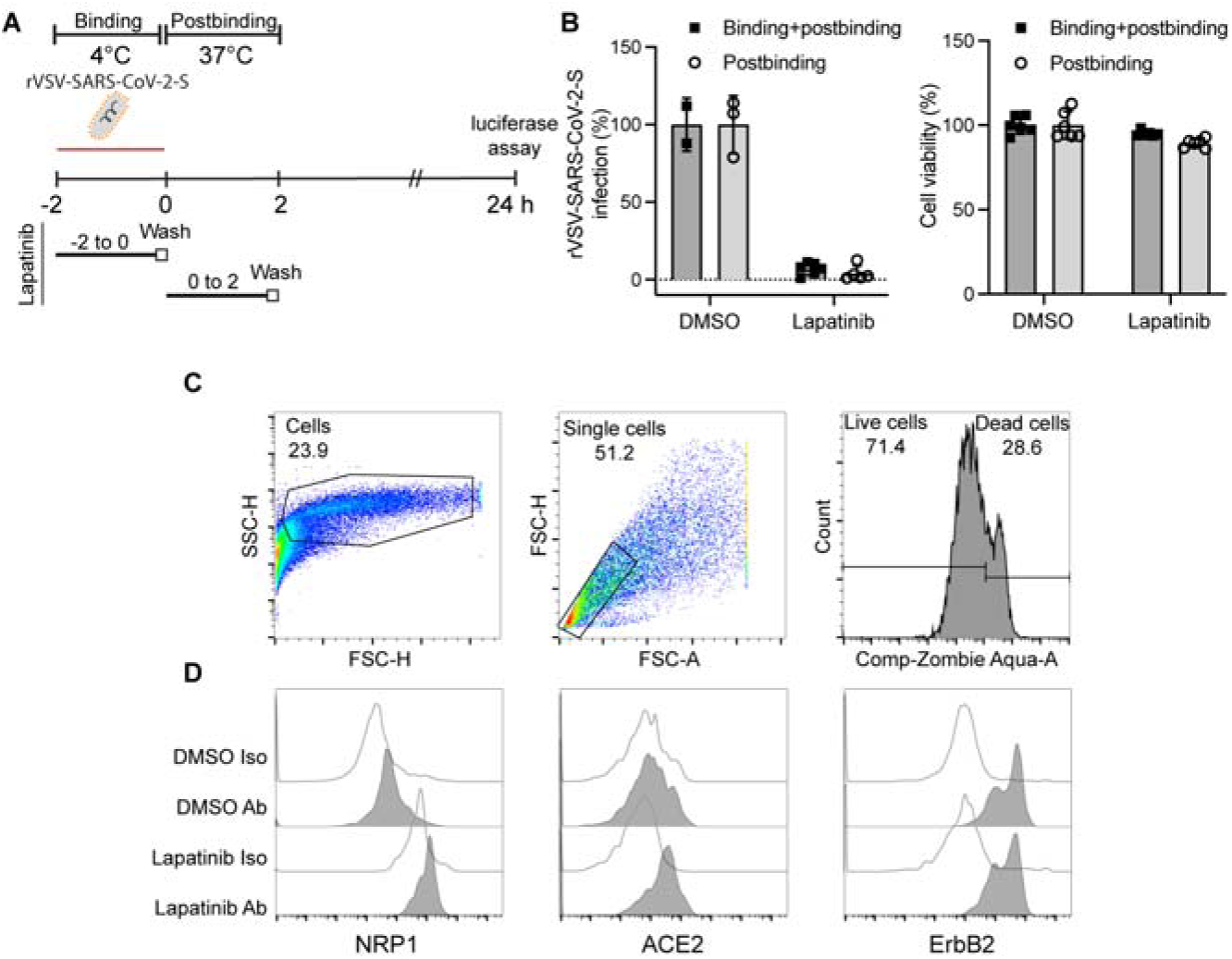
Lapatinib treatment suppresses viral entry at a postbinding stage. Related to figure 5. **A**, Schematic of the temperature-shift experiments shown in panel B**. B,** Vero cells were infected with VSV-SARS-CoV-2-S for 2 hours at 4°C in the presence or absence of 10 µM lapatinib or DMSO before the temperature was shifted to 37°C to initiate infection. 24 hpi virus infection was measured via luciferase assay and cell viability by alamarBlue assay. Values are shown relative to DMSO control. **C**, Representative dot plots showing gating strategy. Cell debris were excluded by size, and dead cells were excluded using Zombie Aqua live/dead staining. **D**, Representative histograms depicting surface expression of NRP1, ACE2 and ErbB2 in SARS-CoV-2-infected and DMSO- or lapatinib-treated cells. Antibody isotype control histograms are also shown.

**Supplemental Figure 7:**
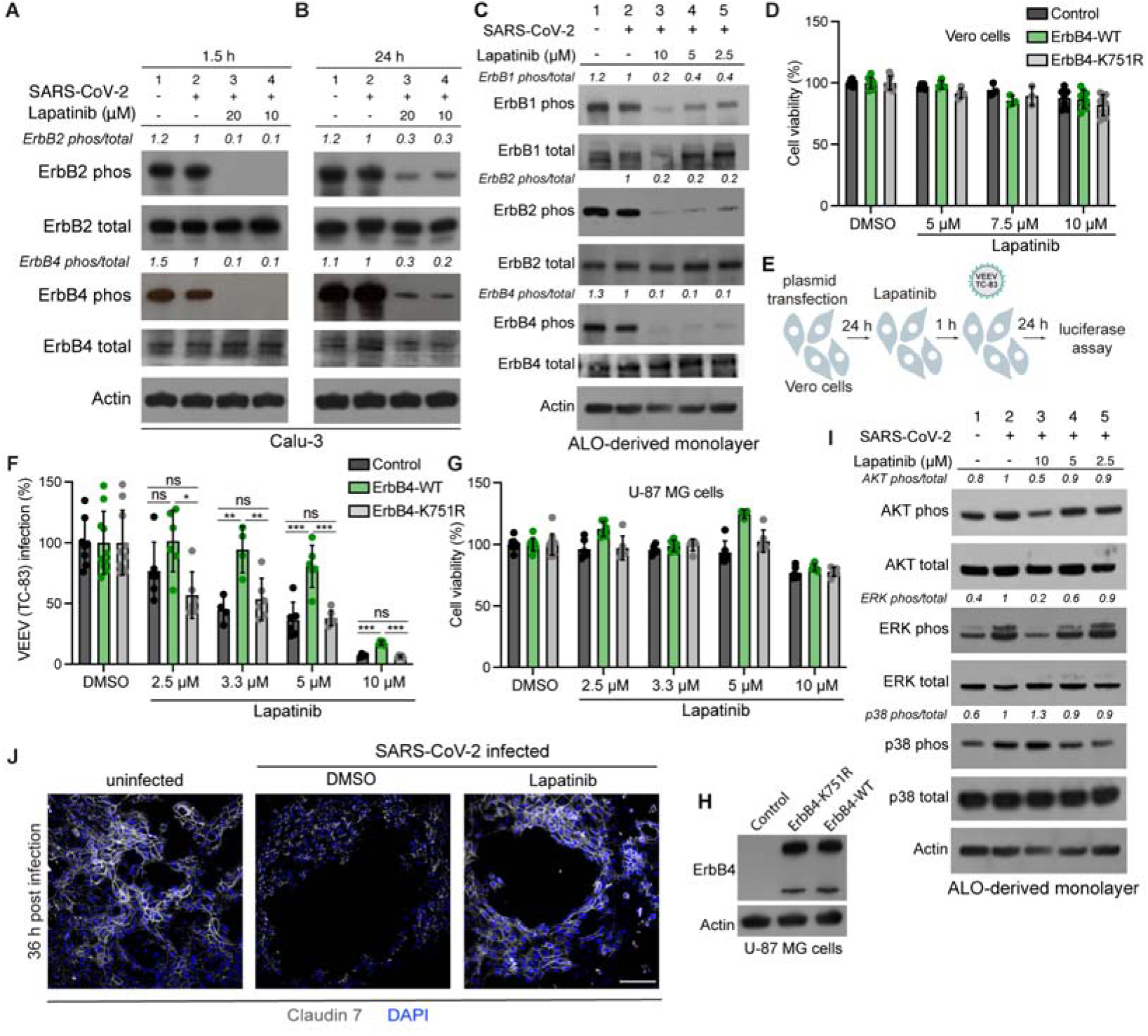
Lapatinib treatment modulates ErbBs and suppresses activation of downstream tissue injury signals. Related to figure 6. **A, B,** ErbB2 and ErbB4 phosphorylation in Calu-3 cells that are uninfected (lane 1), infected and treated with DMSO (lane 2) or infected and treated with lapatinib (lanes 3 and 4) measured by Western blotting 1.5 (**A**) and 24 (**B**) hpi with SARS-CoV-2 (USA-WA1/2020 strain, MOI=1). **C,** Dose-dependent effect of lapatinib treatment on ErbB1, 2 and 4 phosphorylation in ALO-derived monolayers that are uninfected (lane 1), SARS-CoV-2-infected and treated with DMSO (lane 2) or infected and treated with lapatinib (lanes 3-5) measured by Western blotting 48 hpi. Shown are representative membranes blotted for phospho- and total ErbB2, ErbB4, and actin and quantitative phospho- to total ErbB ratio data relative to infected cells treated with DMSO (lane 2). **D,** Vero cell viability measured by alamarBlue assays 48 hours post-transfection of the indicated plasmids. Data relative to the respective DMSO controls are shown. **E,** Schematic of the experiments shown in F-H. **F,** Rescue of VEEV (TC-83) infection in the presence of lapatinib upon ectopic expression of the indicated plasmids measured by luciferase assays 24 hpi in U-87 MG cells. **G,** U-87 MG cell viability measured by alamarBlue assays 48 hours post-transfection of the indicated plasmids. Data relative to the respective DMSO controls are shown**. H**, Level of ErbB4 and actin expression measured via Western blot following transfection of U-87 MG cells with control or ErbB4-expressing plasmids. **I**, Dose-dependent effect of lapatinib treatment on AKT, ERK and p38 MAPK phosphorylation in ALO-derived monolayers that are uninfected (lane 1), SARS-CoV-2-infected and treated with DMSO (lane 2) or infected and treated with lapatinib (lanes 3-5) measured by Western blotting 48 hpi. **J,** Confocal IF microscopy images of Claudin 7 (grey) and DAPI (blue) in naïve or SARS-CoV-2-infected ALO-derived monolayers treated at 4 hpi either with DMSO or 10 μM lapatinib and imaged at 36 hpi. Representative merged images at 20x magnification are shown. Scale bar is 100 µm.

**Supplemental text 1. The HTS of compound libraries for SARS-CoV-2 inhibitors is robust and specific.**

The libraries were screened in two independent experiments. Data were normalized to the median of each plate. The average percent fluorescent area for control wells included in each plate was 102.9±5% for uninfected cells (cell control), 0.1±0.2% for infected untreated cells (virus control), and 0.0±0.164% for infected cells treated with DMSO (**Figure 1C**). The *Z*-score was calculated on the basis of the log2(fold change) (log2FC) with the average and standard deviation of each plate. The Z’ and RZ’ values of each of the 29 screen plates, calculated based on the virus control and cell control wells, were greater than 0.78 and the signal-to-background (S/B) value, representing the ratio of the median value of the raw data between the virus control and the cell control, was greater than 120 (**Supplemental Figure 1B**). The two replicate screens demonstrated good correlation (r= 0.76) (**Supplemental Figure 1C**). Remdesivir and its major metabolite, GS-441524, used as positive controls, demonstrated dose-dependent anti-SARS-CoV-2 activity in this assay (**Supplemental Figure 1D**). Overall, these data indicate that this antiviral assay is robust for HTS and is specific. 40 compounds from the screen were selected according to the cutoff of fluorescence % area greater than 15 in at least one of the two screens, which is 15 times greater than the values obtained with untreated or DMSO treated cells.

**Supplemental text 2. Broad-spectrum antiviral activity of hits.**

The effect of the 18 emerging hit compounds on replication of two unrelated RNA viruses, TC-83, the vaccine strain of the alphavirus VEEV, and the flavivirus, dengue (DENV2) was measured in human astrocytes (U-87 MG) and human hepatoma (Huh7) cells, respectively, both via luciferase assays. Lycorine, calcimycin, monensin, azaserine, gedunin, and the kinase inhibitors lapatinib and AG 879 dose-dependently inhibited replication of TC-83 and DENV2 in addition to SARS-CoV-2 (**Supplemental Figure 3A, B**). Several compounds, such as tyrphostin A9, an investigational inhibitor of PDGFR (platelet-derived growth factor receptor), and fluspirilene, a neuroleptic agent, demonstrated more potent anti-TC-83 and DENV2 activity than anti-SARS-CoV-2 activity, and others showed variable activity against one or two of these viruses. Salbutamol demonstrated minimal to no activity against all three viruses (**Supplemental Figure 3A, B**).

**Supplemental text 3. Safety considerations and drug-drug interactions of lapatinib. Related to discussion.**

Although toxicity is a concern when targeting host functions, lapatinib has a favorable safety profile, particularly when used as a monotherapy and for short durations, as those required to treat acute infections.

Notably, lapatinib’s safety profile in the package insert was based on data from over 12,000 patients with advanced cancer who received lapatinib in combination with capecitabine or trastuzumab plus an aromatase inhibitor and for long durations(1–3). As monotherapy, lapatinib was tested in several open-label studies with a median duration of 7-28 weeks in patients with advanced cancer(4–11). The most common adverse events attributed to lapatinib were diarrhea, rash, nausea, pruritus, and fatigue, with diarrhea being the most common adverse event resulting in drug discontinuation. The most common laboratory abnormalities with combination therapy were increased liver function tests, which were infrequently severe(1–3, 12). More severe adverse events including transient, reversible decreases in left ventricular ejection fraction, prolongation of QT interval, and hepatotoxicity, were also documented, yet infrequently, and with the exception of cardiac toxicity, primarily in patients receiving lapatinib in combination treatment(1, 10, 13, 14).

Notably, unlike erlotinib and gefitinib, lapatinib monotherapy has not been associated with pneumonitis, interstitial lung disease or lung fibrosis(4–11). The estimated incidence of 0.2% for these adverse effects is based on patients receiving lapatinib in combination with other drugs(15–20) known to cause pneumonitis and/or lung fibrosis(21–23), and sometimes also with radiation, for a median duration of 24-45 weeks. We predict that lapatinib’s distinct off-target profile accounts for this difference in the occurrence of these adverse events. Indeed, cyclin G-associated kinase (GAK), an off-target of erlotinib (K*_D_*=3.1 nM, IC_50_=0.88 µM) and gefitinib (K*_D_*=6.5 nM, IC_50_=0.41 µM), but not of lapatinib (K*_D_*=980 nM, IC_50_>5 µM)(24), has been implicated in pulmonary alveolar function and stem cell regeneration, and its inhibition is thought to be the mechanism underlying gefinitib- and erlotinib-induced lung toxicity(25, 26).

An important consideration with lapatinib is, however, its potential for drug-drug interactions. Since metabolized by CYP3A4, concurrent use of suppressors of CYP3A4 should be avoided to reduce risk of QT prolongation. Concurrent treatment with CYP3A4 inducers should also be avoided, as this can reduce lapatinib’s levels to sub-therapeutic. Of particular relevance is the CYP3A4 inducer dexamethasone used as standard of care for moderate COVID-19 patients.

Since other steroids do not induce CYP3A4, lapatinib could be studied in combination with hydrocortisone or prednisone, which have been shown to comparably protect COVID-19 patients(27–29).

**Supplemental text 4. Broad-spectrum potential of other hits emerging in the screen. Related to discussion.**

Another approved anticancer drug that emerged in the HTS was sunitinib, a multi-kinase inhibitor that we have shown to protect mice from DENV and EBOV challenges when given in combination with erlotinib by inhibiting NAK-mediated intracellular viral trafficking(30–32). Sunitinib was recently shown by others to suppress pan-corona pseudotyped viral infections(33) and by us to suppress WT SARS-CoV-2 infection(34). AG 879, another kinase inhibitor demonstrating anti-SARS-CoV-2 activity, was reported to suppress replication of multiple viruses including a mouse hepatitis virus (Coronaviridae) in cultured cells and to protect mice from influenza A virus (IAV) challenge(35–37). Nevertheless, since we could not confirm its anti-ErbB activity, the precise target(s) mediating the antiviral effect remain to be elucidated.

Ion transport across cell membranes is another function that emerged in our HTS as a candidate target for anti-SARS-CoV-2 approaches. Among the hits was tetrandine, a calcium channel blocker with anti-inflammatory and anti-fibrogenic properties used as a medicinal herb for the treatment of lung silicosis, liver cirrhosis, and rheumatoid arthritis(38). Tetrandine was previously shown to inhibit EBOV entry in cultured cells and protect EBOV-infected mice by inhibiting endosomal calcium channels(39). Monensin, an antiprotozoal agent, and calcimycin, shown to inhibit VSV and IAV infections(40, 41), are both ionophores that facilitate the transport of sodium/potassium and calcium across the membrane, respectively. Spiperone, an activator of chloride channels licensed in Japan for the treatment of schizophrenia, was another hit.

The emergence of gedunin, a natural product that inhibits HSP90 and has anti-inflammatory properties, suggests a potential role for HSP90 in SARS-CoV-2 infection, as in other viral infections(42, 43). Lycorine, a protein synthesis inhibitor(44) was also shown to suppress replication of multiple viruses including SARS-CoV in cultured cells(45–48) and mortality of mice infected with human enterovirus 71(49). The underlying mechanism of action in influenza was thought to be inhibition of export of viral ribonucleoprotein complexes from the nucleus(45), yet lycorine also exhibits anti-inflammatory effects(50). Azaserine is a natural serine derivative that irreversibly inhibits γ-glutamyltransferase in the metabolic hexosamine pathway. Independently of this target, it was shown to protect from endothelial cell inflammation and injury(51).

Aurothioglucose has been used for the treatment of rheumatoid arthritis and is thought to inhibit the activity of adenylyl cyclase in inflammatory pathways(52). Ac-Leu-Leu-Nle-CHO is used as a research tool to inhibit calpain 1 and 2 (CAPN1 and 2)(53), cysteine proteases required for SARS-CoV(54), echovirus 1(55) and herpes simplex virus(56) infections. Targeting calpain proteases was shown to inhibit SARS-CoV-2(57), SARS-CoV(58) and IAV replication(59) and to exert anti-inflammatory and tissue protective effects(60, 61) including in a reovirus-induced myocarditis mouse model(62). Beyond their host-targeted effects, Ac-Leu-Leu-Nle-CHO and aurothioglucose may have direct antiviral effects against the SARS-CoV-2 M^pro^ or 3C-like proteases, respectively(57, 63). Lastly, josamycin is a natural macrolide antibiotic with an anti-inflammatory activity used in humans in Europe and Japan. Other macrolides have shown anti-IAV and anti-inflammatory activities(64). These findings reveal candidate targets for anti-SARS-CoV-2 approaches. Moreover, they underscore the potential utility of natural products as broad-spectrum antivirals, yet limited scalability typically challenges the use of these products.

### Supplemental methods

#### Compounds

The Microsource Spectrum, two Biomol and LOPAC libraries were available at the Stanford High-Throughput Bioscience Center. Small molecule inhibitors were purchased from MedchemExpress or Cayman Chemical. Dinaciclib and ribociclib were a gift from Dr. Mardo Koivomagi.

#### Plasmids

Plasmids used for production of SARS-CoV-2 pseudovirus were a gift from Jing Lin (Vitalant, San Francisco). The rSARS-CoV-2/WT and rSARS-CoV-2/Nluc (rSARS-CoV-2 expressing Nluc-reporter gene) plasmids were a gift from <u>Luis Martinez-Sobrido</u> and were generated as previously described (65, 66). Flag-tagged SARS-CoV-2 (2019-nCoV) Spike S1 expression plasmid was purchased from Sino Biological (#VG40591-CF). Plasmid encoding VEEV TC-83 with a nanoluciferase reporter (VEEV TC-83-Cap-nLuc-Tav) was a gift from Dr. William B. Klimstra (Department of Immunology, University of Pittsburgh) (67). DENV2 (New Guinea C strain) TSV01 Renilla reporter plasmid (pACYC NGC FL) was a gift from Pei-Yong Shi (University of Texas Medical Branch) (68). pDONR223-EGFR, pDONR223-ErbB2, pDONR223-ErbB4 were a gift from William Hahn & David Root (Addgene)(69). ORFs were recombined into a gateway-compatible pGluc destination vector using Gateway technology (Invitrogen). Mutations were introduced by site-directed mutagenesis using the QuikChange Lightning Site-Directed Mutagenesis Kit (Agilent).

#### Cells

The African green monkey kidney cell line (Vero E6) constitutively expressing enhanced green fluorescent protein (eGFP) was provided by Dr. Marnix Van Loock (Janssen Pharmaceutica, Beerse, Belgium) (70). Cells were maintained in Dulbecco’s modified Eagle’s medium (DMEM, Gibco) supplemented with 10% v/v fetal calf serum (Biowest), 0.075% sodium bicarbonate and 1x Pen-strep (Gibco). Vero E6, Vero, Calu-3, HEK-293T, U-87 MG, and BHK-21 cells (ATCC) were maintained in DMEM supplemented with 10% fetal bovine serum (FBS, Omega Scientific, Inc), 1% L-glutamine 200mM, 1% penicillin-streptomycin, 1% nonessential amino acids, 1% HEPES (Gibco), 1% Sodium pyruvate (Thermofisher scientific). A549-NRP1^KO^ cells were grown in DMEM:Hams F12 (Cytiva) supplemented with 5% FBS. TMPRSS2-expressing Vero E6 cells were maintained in DMEM supplemented with 10% FBS and G418 (1 mg/mL) (Thermofisher, Gibco). All cells were maintained in a humidified incubator with 5% CO_2_ at 37°C and tested negative for mycoplasma by MycoAlert (Lonza, Morristown, NJ).

## References

1. Wu Z, and McGoogan JM. Characteristics of and Important Lessons From the Coronavirus Disease 2019 (COVID-19) Outbreak in China: Summary of a Report of 72c:314 Cases From the Chinese Center for Disease Control and Prevention. JAMA. 2020;323(13):1239–42.

2. Spagnolo P, Balestro E, Aliberti S, Cocconcelli E, Biondini D, Casa GD, et al. Pulmonary fibrosis secondary to COVID-19: a call to arms? The Lancet Respiratory medicine. 2020;8(8):750–2.

3. Weaver SC, Ferro C, Barrera R, Boshell J, and Navarro JC. Venezuelan equine encephalitis. Annual review of entomology. 2004;49:141–74.

4. Leffel EK, and Reed DS. Marburg and Ebola viruses as aerosol threats. Biosecurity and bioterrorism : biodefense strategy, practice, and science. 2004;2(3):186–91.

5. Bekerman E, and Einav S. Combating emerging viral threats. Science. 2015;348(6232):282–3.

6. Szemiel AM, Merits A, Orton RJ, MacLean OA, Pinto RM, Wickenhagen A, et al. In vitro selection of Remdesivir resistance suggests evolutionary predictability of SARS-CoV-2. PLoS pathogens. 2021;17(9):e1009929.

7. Iketani S, Mohri H, Culbertson B, Hong SJ, Duan Y, Luck MI, et al. Multiple pathways for SARS-CoV-2 resistance to nirmatrelvir. Nature. 2023;613(7944):558–64.

8. Pandit JA, Radin JM, Chiang D, Spencer E, Pawelek J, Diwan M, et al. The Paxlovid Rebound Study: A Prospective Cohort Study to Evaluate Viral and Symptom Rebound Differences Between Paxlovid and Untreated COVID-19 Participants. medRxiv. 2022:2022.11.14.22282195.

9. Bekerman E, Neveu G, Shulla A, Brannan J, Pu S-Y, Wang S, et al. Anticancer kinase inhibitors impair intracellular viral trafficking and exert broad-spectrum antiviral effects. The Journal of clinical investigation. 2017;127(4).

10. Ivens T, Van den Eynde C, Van Acker K, Nijs E, Dams G, Bettens E, et al. Development of a homogeneous screening assay for automated detection of antiviral agents active against severe acute respiratory syndrome-associated coronavirus. J Virol Methods. 2005;129(1):56–63.

11. Saul S, and Einav S. Old Drugs for a New Virus: Repurposed Approaches for Combating COVID-19. ACS infectious diseases. 2020.

12. Levitzki A, and Gazit A. Tyrosine kinase inhibition: an approach to drug development. Science. 1995;267(5205):1782–8.

13. Tindle C, Fuller M, Fonseca A, Taheri S, Ibeawuchi SR, Beutler N, et al. Adult Stem Cell-derived Complete Lung Organoid Models Emulate Lung Disease in COVID-19. bioRxiv : the preprint server for biology. 2020.

14. Prichard MN, and Shipman C. Analysis of combinations of antiviral drugs and design of effective multidrug therapies. Antiviral therapy. 1996;1(1):9–20.

15. Chung D, Schroeder CE, Sotsky J, Yao T, Roy S, Smith RA, et al. Probe Reports from the NIH Molecular Libraries Program. Bethesda (MD): National Center for Biotechnology Information (US); 2010.

16. Schäfer A, Brooke CB, Whitmore AC, and Johnston RE. The role of the blood-brain barrier during Venezuelan equine encephalitis virus infection. J Virol. 2011;85(20):10682–90.

17. David K. Schaffer JPW, Ronald S. Reiserer, Michael D. Geuy, Eric C. Spivey, Clayton M. Britt, Jacquelyn A. Brown, Dmitry A. Markov, Shannon Faley, Lisa J. Mccawley, Philip C. Samson. USA; 2021.

18. Boghdeh NA, Risner KH, Barrera MD, Britt CM, Schaffer DK, Alem F, et al. Application of a Human Blood Brain Barrier Organ-on-a-Chip Model to Evaluate Small Molecule Effectiveness against Venezuelan Equine Encephalitis Virus. Viruses. 2022;14(12).

19. Nair AB, and Jacob S. A simple practice guide for dose conversion between animals and human. Journal of basic and clinical pharmacy. 2016;7(2):27–31.

20. Spector NL, Robertson FC, Bacus S, Blackwell K, Smith DA, Glenn K, et al. Lapatinib Plasma and Tumor Concentrations and Effects on HER Receptor Phosphorylation in Tumor. PLOS ONE. 2015;10(11):e0142845.

21. European Medicines Agency Evaluation of Medicines for Human Use; 2008.

22. Finigan JH, Downey GP, and Kern JA. Human epidermal growth factor receptor signaling in acute lung injury. American journal of respiratory cell and molecular biology. 2012;47(4):395–404.

23. Chen J, Kinoshita T, Sukbuntherng J, Chang BY, and Elias L. Ibrutinib Inhibits ERBB Receptor Tyrosine Kinases and HER2-Amplified Breast Cancer Cell Growth. Mol Cancer Ther. 2016;15(12):2835–44.

24. Qiu C, Tarrant MK, Choi SH, Sathyamurthy A, Bose R, Banjade S, et al. Mechanism of activation and inhibition of the HER4/ErbB4 kinase. Structure (London, England : 1993). 2008;16(3):460–7.

25. Hickinson DM, Klinowska T, Speake G, Vincent J, Trigwell C, Anderton J, et al. AZD8931, an Equipotent, Reversible Inhibitor of Signaling by Epidermal Growth Factor Receptor, ERBB2 (HER2), and ERBB3: A Unique Agent for Simultaneous ERBB Receptor Blockade in Cancer. Clinical Cancer Research. 2010;16(4):1159–69.

26. Jackson CB, Farzan M, Chen B, and Choe H. Mechanisms of SARS-CoV-2 entry into cells. Nature Reviews Molecular Cell Biology. 2022;23(1):3–20.

27. Hoffmann M, Kleine-Weber H, Schroeder S, Krüger N, Herrler T, Erichsen S, et al. SARS-CoV-2 Cell Entry Depends on ACE2 and TMPRSS2 and Is Blocked by a Clinically Proven Protease Inhibitor. Cell. 2020;181(2):271–80.e8.

28. Daly JL, Simonetti B, Klein K, Chen K-E, Williamson MK, Antón-Plágaro C, et al. Neuropilin-1 is a host factor for SARS-CoV-2 infection. Science. 2020;370(6518):861–5.

29. Sánchez-Martín M, and Pandiella A. Differential action of small molecule HER kinase inhibitors on receptor heterodimerization: therapeutic implications. International journal of cancer. 2012;131(1):244–52.

30. Portales AE, Mustafá ER, McCarthy CI, Cornejo MP, Couto PM, Gironacci MM, et al. ACE2 internalization induced by a SARS-CoV-2 recombinant protein is modulated by angiotensin II type 1 and bradykinin 2 receptors. Life sciences. 2022;293:120284.

31. Ogunlade BO, Lazartigues E, and Filipeanu CM. Angiotensin Type 1 Receptor-Dependent Internalization of SARS-CoV-2 by Angiotensin-Converting Enzyme 2. Hypertension. 2021;77(4):e42–e3.

32. Ma X, Yu X, and Zhou Q. The IL1β-HER2-CLDN18/CLDN4 axis mediates lung barrier damage in ARDS. Aging (Albany NY). 2020;12(4):3249–65.

33. Faress JA, Nethery DE, Kern EF, Eisenberg R, Jacono FJ, Allen CL, et al. Bleomycin-induced pulmonary fibrosis is attenuated by a monoclonal antibody targeting HER2. Journal of applied physiology (Bethesda, Md : 1985). 2007;103(6):2077–83.

34. Hardie WD, Kerlakian CB, Bruno MD, Huelsman KM, Wert SE, Glasser SW, et al. Reversal of lung lesions in transgenic transforming growth factor alpha mice by expression of mutant epidermal growth factor receptor. American journal of respiratory cell and molecular biology. 1996;15(4):499–508.

35. Li R, Zou X, Huang H, Yu Y, Zhang H, Liu P, et al. HMGB1/PI3K/Akt/mTOR Signaling Participates in the Pathological Process of Acute Lung Injury by Regulating the Maturation and Function of Dendritic Cells. Frontiers in Immunology. 2020;11(1104).

36. Appelberg S, Gupta S, Svensson Akusjärvi S, Ambikan AT, Mikaeloff F, Saccon E, et al. Dysregulation in Akt/mTOR/HIF-1 signaling identified by proteo-transcriptomics of SARS-CoV-2 infected cells. Emerging Microbes & Infections. 2020;9(1):1748–60.

37. Bouhaddou M, Memon D, Meyer B, White KM, Rezelj VV, Correa Marrero M, et al. The Global Phosphorylation Landscape of SARS-CoV-2 Infection. Cell. 2020;182(3):685–712.e19.

38. Del Valle DM, Kim-Schulze S, Huang H-H, Beckmann ND, Nirenberg S, Wang B, et al. An inflammatory cytokine signature predicts COVID-19 severity and survival. Nature medicine. 2020;26(10):1636–43.

39. Sokol CL, and Luster AD. The chemokine system in innate immunity. Cold Spring Harbor perspectives in biology. 2015;7(5).

40. Bierman A, Yerrapureddy A, Reddy NM, Hassoun PM, and Reddy SP. Epidermal growth factor receptor (EGFR) regulates mechanical ventilation-induced lung injury in mice. Transl Res. 2008;152(6):265–72.

41. Ishii Y, Fujimoto S, and Fukuda T. Gefitinib prevents bleomycin-induced lung fibrosis in mice. American journal of respiratory and critical care medicine. 2006;174(5):550–6.

42. Hardie WD, Davidson C, Ikegami M, Leikauf GD, Cras TDL, Prestridge A, et al. EGF receptor tyrosine kinase inhibitors diminish transforming growth factor-α-induced pulmonary fibrosis. American Journal of Physiology-Lung Cellular and Molecular Physiology. 2008;294(6):L1217–L25.

43. Ho J, Moyes DL, Tavassoli M, and Naglik JR. The Role of ErbB Receptors in Infection. Trends in microbiology. 2017;25(11):942–52.

44. Drayman N, DeMarco JK, Jones KA, Azizi SA, Froggatt HM, Tan K, et al. Masitinib is a broad coronavirus 3CL inhibitor that blocks replication of SARS-CoV-2. Science. 2021;373(6557):931–6.

45. Mizutani T, Fukushi S, Saijo M, Kurane I, and Morikawa S. Phosphorylation of p38 MAPK and its downstream targets in SARS coronavirus-infected cells. Biochem Biophys Res Commun. 2004;319(4):1228–34.

46. Kindrachuk J, Ork B, Hart BJ, Mazur S, Holbrook MR, Frieman MB, et al. Antiviral Potential of ERK/MAPK and PI3K/AKT/mTOR Signaling Modulation for Middle East Respiratory Syndrome Coronavirus Infection as Identified by Temporal Kinome Analysis. Antimicrobial agents and chemotherapy. 2015;59(2):1088–99.

47. Venkataraman T, Coleman CM, and Frieman MB. Overactive Epidermal Growth Factor Receptor Signaling Leads to Increased Fibrosis after Severe Acute Respiratory Syndrome Coronavirus Infection. Journal of virology. 2017;91(12):e00182–17.

48. Li S-W, Wang C-Y, Jou Y-J, Yang T-C, Huang S-H, Wan L, et al. SARS coronavirus papain-like protease induces Egr-1-dependent up-regulation of TGF-β1 via ROS/p38 MAPK/STAT3 pathway. Scientific Reports. 2016;6(1):25754.

49. Simões e Silva AC, Silveira KD, Ferreira AJ, and Teixeira MM. ACE2, angiotensin-(1-7) and Mas receptor axis in inflammation and fibrosis. Br J Pharmacol. 2013;169(3):477–92.

50. Akhtar S, Chandrasekhar B, Attur S, Dhaunsi GS, Yousif MHM, and Benter IF. Transactivation of ErbB Family of Receptor Tyrosine Kinases Is Inhibited by Angiotensin-(1-7) via Its Mas Receptor. PloS one. 2015;10(11):e0141657-e.

51. Freeman MC, Peek CT, Becker MM, Smith EC, and Denison MR. Coronaviruses Induce Entry-Independent, Continuous Macropinocytosis. mBio. 2014;5(4):e01340–14.

52. Treon SP, Castillo JJ, Skarbnik AP, Soumerai JD, Ghobrial IM, Guerrera ML, et al. The BTK inhibitor ibrutinib may protect against pulmonary injury in COVID-19-infected patients. Blood. 2020;135(21):1912–5.

53. Knight ZA, Lin H, and Shokat KM. Targeting the cancer kinome through polypharmacology. Nat Rev Cancer. 2010;10(2):130–7.

54. Hudachek SF, and Gustafson DL. Physiologically based pharmacokinetic model of lapatinib developed in mice and scaled to humans. Journal of pharmacokinetics and pharmacodynamics. 2013;40(2):157–76.

55. Ivens T, Eynde CVd, Acker KV, Nijs E, Dams G, Bettens E, et al. Development of a homogeneous screening assay for automated detection of antiviral agents active against severe acute respiratory syndrome-associated coronavirus. Journal of virological methods. 2005;129(1):56–63.

56. Spiteri G, Fielding J, Diercke M, Campese C, Enouf V, Gaymard A, et al. First cases of coronavirus disease 2019 (COVID-19) in the WHO European Region, 24 January to 21 February 2020. Euro Surveill. 2020;25(9).

57. Chiem K, Morales Vasquez D, Park J-G, Platt RN, Anderson T, Walter MR, et al. Generation and Characterization of Recombinant SARS-CoV-2 Expressing Reporter Genes. Journal of virology. 2021;95(7):e02209–20.

58. Karim M, Saul S, Ghita L, Sahoo MK, Ye C, Bhalla N, et al. Numb-associated kinases are required for SARS-CoV-2 infection and are cellular targets for antiviral strategies. Antiviral Res. 2022;204:105367.

59. Boudewijns R, Thibaut HJ, Kaptein SJF, Li R, Vergote V, Seldeslachts L, et al. STAT2 signaling as double-edged sword restricting viral dissemination but driving severe pneumonia in SARS-CoV-2 infected hamsters. bioRxiv. 2020:2020.04.23.056838.

60. Prichard MN, and Shipman C. A three-dimensional model to analyze drug-drug interactions. Antiviral Res. 1990;14(4-5):181–205.

## Supplemental references

1. Novartis. Tykerb (U.S. package insert). 2018.

2. Pivot X, Manikhas A, Żurawski B, Chmielowska E, Karaszewska B, Allerton R, et al. CEREBEL (EGF111438): A Phase III, Randomized, Open-Label Study of Lapatinib Plus Capecitabine Versus Trastuzumab Plus Capecitabine in Patients With Human Epidermal Growth Factor Receptor 2-Positive Metastatic Breast Cancer. J Clin Oncol. 2015;33(14):1564–73.

3. Schwartzberg LS, Franco SX, Florance A, O’Rourke L, Maltzman J, and Johnston S. Lapatinib plus letrozole as first-line therapy for HER-2+ hormone receptor-positive metastatic breast cancer. The oncologist. 2010;15(2):122–9.

4. Blackwell KL, Pegram MD, Tan-Chiu E, Schwartzberg LS, Arbushites MC, Maltzman JD, et al. Single-agent lapatinib for HER2-overexpressing advanced or metastatic breast cancer that progressed on first- or second-line trastuzumab-containing regimens. Ann Oncol. 2009;20(6):1026–31.

5. Blackwell KL, Burstein HJ, Storniolo AM, Rugo H, Sledge G, Koehler M, et al. Randomized study of Lapatinib alone or in combination with trastuzumab in women with ErbB2-positive, trastuzumab-refractory metastatic breast cancer. J Clin Oncol. 2010;28(7):1124–30.

6. Burris HA, 3rd, Hurwitz HI, Dees EC, Dowlati A, Blackwell KL, O’Neil B, et al. Phase I safety, pharmacokinetics, and clinical activity study of lapatinib (GW572016), a reversible dual inhibitor of epidermal growth factor receptor tyrosine kinases, in heavily pretreated patients with metastatic carcinomas. J Clin Oncol. 2005;23(23):5305–13.

7. Burstein HJ, Storniolo AM, Franco S, Forster J, Stein S, Rubin S, et al. A phase II study of lapatinib monotherapy in chemotherapy-refractory HER2-positive and HER2-negative advanced or metastatic breast cancer. Ann Oncol. 2008;19(6):1068–74.

8. Gomez HL, Doval DC, Chavez MA, Ang PC-S, Aziz Z, Nag S, et al. Efficacy and Safety of Lapatinib As First-Line Therapy for ErbB2-Amplified Locally Advanced or Metastatic Breast Cancer. Journal of Clinical Oncology. 2008;26(18):2999–3005.

9. Hurvitz SA, and Kakkar R. Role of lapatinib alone or in combination in the treatment of HER2-positive breast cancer. Breast Cancer (Dove Med Press). 2012;4:35–51.

10. Perez EA, Koehler M, Byrne J, Preston AJ, Rappold E, and Ewer MS. Cardiac safety of lapatinib: pooled analysis of 3689 patients enrolled in clinical trials. Mayo Clinic proceedings. 2008;83(6):679–86.

11. Toi M, Iwata H, Fujiwara Y, Ito Y, Nakamura S, Tokuda Y, et al. Lapatinib monotherapy in patients with relapsed, advanced, or metastatic breast cancer: efficacy, safety, and biomarker results from Japanese patients phase II studies. British journal of cancer. 2009;101(10):1676–82.

12. Piccart-Gebhart M, Holmes E, Baselga J, de Azambuja E, Dueck AC, Viale G, et al. Adjuvant Lapatinib and Trastuzumab for Early Human Epidermal Growth Factor Receptor 2-Positive Breast Cancer: Results From the Randomized Phase III Adjuvant Lapatinib and/or Trastuzumab Treatment Optimization Trial. J Clin Oncol. 2016;34(10):1034–42.

13. Kloth JSL, Pagani A, Verboom MC, Malovini A, Napolitano C, Kruit WHJ, et al. Incidence and relevance of QTc-interval prolongation caused by tyrosine kinase inhibitors. British journal of cancer. 2015;112(6):1011–6.

14. Dogan E, Yorgun H, Petekkaya I, Ozer N, Altundag K, and Ozisik Y. Evaluation of cardiac safety of lapatinib therapy for ErbB2-positive metastatic breast cancer: a single center experience. Medical oncology (Northwood, London, England). 2012;29(5):3232–9.

15. Hackshaw MD, Danysh HE, Singh J, Ritchey ME, Ladner A, Taitt C, et al. Incidence of pneumonitis/interstitial lung disease induced by HER2-targeting therapy for HER2-positive metastatic breast cancer. Breast cancer research and treatment. 2020;183(1):23–39.

16. Jagiello-Gruszfeld A, Tjulandin S, Dobrovolskaya N, Manikhas A, Pienkowski T, DeSilvio M, et al. A single-arm phase II trial of first-line paclitaxel in combination with lapatinib in HER2-overexpressing metastatic breast cancer. Oncology. 2010;79(1-2):129–35.

17. Capri G, Chang J, Chen SC, Conte P, Cwiertka K, Jerusalem G, et al. An open-label expanded access study of lapatinib and capecitabine in patients with HER2-overexpressing locally advanced or metastatic breast cancer. Ann Oncol. 2010;21(3):474–80.

18. Xu B-H, Jiang Z-F, Chua D, Shao Z-M, Luo R-C, Wang X-J, et al. Lapatinib plus capecitabine in treating HER2-positive advanced breast cancer: efficacy, safety, and biomarker results from Chinese patients. Chin J Cancer. 2011;30(5):327–35.

19. 19. Bates CA, Zhao B, Schlobohm A, Asquith C, Zuercher W, and Barkauskas C. 2019.

20. Brenner T, Motsch J, Werner J, Grenacher L, Martin E, and Hofer S. Rapid-Onset Acute Respiratory Distress Syndrome (ARDS) in a Patient Undergoing Metastatic Liver Resection: A Case Report and Review of the Literature. Anesthesiol Res Pract. 2010;2010:586425.

21. Chan AK, Choo BA, and Glaholm J. Pulmonary toxicity with oxaliplatin and capecitabine/5-Fluorouracil chemotherapy: a case report and review of the literature. Onkologie. 2011;34(8-9):443–6.

22. Torrisi JM, Schwartz LH, Gollub MJ, Ginsberg MS, Bosl GJ, and Hricak H. CT findings of chemotherapy-induced toxicity: what radiologists need to know about the clinical and radiologic manifestations of chemotherapy toxicity. Radiology. 2011;258(1):41–56.

23. Bielopolski D, Evron E, Moreh-Rahav O, Landes M, Stemmer SM, and Salamon F. Paclitaxel-induced pneumonitis in patients with breast cancer: case series and review of the literature. Journal of chemotherapy (Florence, Italy). 2017;29(2):113–7.

24. Asquith CRM, Laitinen T, Bennett JM, Wells CI, Elkins JM, Zuercher WJ, et al. Design and Analysis of the 4-Anilinoquin(az)oline Kinase Inhibition Profiles of GAK/SLK/STK10 Using Quantitative Structure-Activity Relationships. ChemMedChem. 2020;15(1):26–49.

25. Tabara H, Naito Y, Ito A, Katsuma A, Sakurai MA, Ohno S, et al. Neonatal lethality in knockout mice expressing the kinase-dead form of the gefitinib target GAK is caused by pulmonary dysfunction. PLoS One. 2011;6(10):e26034.

26. Bates CA, Zhao B, Schlobohm A, Asquith C, Zuercher W, and Barkauskas CE. A61 EPITHELIAL BIOLOGY. American Thoracic Society; 2019:A2125-A.

27. Group TWREAfC-TW. Association Between Administration of Systemic Corticosteroids and Mortality Among Critically Ill Patients With COVID-19: A Meta-analysis. JAMA. 2020;324(13):1330–41.

28. Dequin P-F, Heming N, Meziani F, Plantefève G, Voiriot G, Badié J, et al. Effect of Hydrocortisone on 21-Day Mortality or Respiratory Support Among Critically Ill Patients With COVID-19: A Randomized Clinical Trial. JAMA. 2020;324(13):1298–306.

29. Investigators TWCftR-C. Effect of Hydrocortisone on Mortality and Organ Support in Patients With Severe COVID-19: The REMAP-CAP COVID-19 Corticosteroid Domain Randomized Clinical Trial. JAMA. 2020;324(13):1317–29.

30. Neveu G, Barouch-Bentov R, Ziv-Av A, Gerber D, Jacob Y, and Einav S. Identification and Targeting of an Interaction between a Tyrosine Motif within Hepatitis C Virus Core Protein and AP2M1 Essential for Viral Assembly. PLoS pathogens. 2012;8(8):e1002845.

31. Bekerman E, Neveu G, Shulla A, Brannan J, Pu S-Y, Wang S, et al. Anticancer kinase inhibitors impair intracellular viral trafficking and exert broad-spectrum antiviral effects. The Journal of clinical investigation. 2017;127(4).

32. Pu S, Schor S, Karim M, Saul S, Robinson M, Kumar S, et al. BIKE regulates dengue virus infection and is a cellular target for broad-spectrum antivirals. Antiviral Res. 2020;184:104966.

33. Wang P-G, Tang D-J, Hua Z, Wang Z, and An J. Sunitinib reduces the infection of SARS-CoV, MERS-CoV and SARS-CoV-2 partially by inhibiting AP2M1 phosphorylation. Cell Discovery. 2020;6(1):71.

34. Karim M, Saul S, Ghita L, Sahoo MK, Ye C, Bhalla N, et al. Numb-associated kinases are required for SARS-CoV-2 infection and are cellular targets for antiviral strategies. Antiviral Res. 2022;204:105367.

35. Kumar N, Liang Y, Parslow TG, and Liang Y. Receptor Tyrosine Kinase Inhibitors Block Multiple Steps of Influenza A Virus Replication. Journal of virology. 2011;85(6):2818–27.

36. Kumar N, Sharma NR, Ly H, Parslow TG, and Liang Y. Receptor tyrosine kinase inhibitors that block replication of influenza a and other viruses. Antimicrobial agents and chemotherapy. 2011;55(12):5553–9.

37. Zoeller RA, and Geoghegan-Barek K. A cell-based high-throughput screen identifies tyrphostin AG 879 as an inhibitor of animal cell phospholipid and fatty acid biosynthesis. Biochemistry and Biophysics Reports. 2019;18:100621.

38. Kwan CY, and Achike FI. Tetrandrine and related bis-benzylisoquinoline alkaloids from medicinal herbs: cardiovascular effects and mechanisms of action. Acta pharmacologica Sinica. 2002;23(12):1057–68.

39. Sakurai Y, Kolokoltsov AA, Chen CC, Tidwell MW, Bauta WE, Klugbauer N, et al. Ebola virus. Two-pore channels control Ebola virus host cell entry and are drug targets for disease treatment. Science. 2015;347(6225):995–8.

40. Onishi E, Natori K, and Yamazaki S. The antiviral effect of phorbol ester and calcium ionophore A23187 is not mediated by interferons. Journal of interferon research. 1991;11(3):171–5.

41. Marois I, Cloutier A, Meunier I, Weingartl HM, Cantin AM, and Richter MV. Inhibition of Influenza Virus Replication by Targeting Broad Host Cell Pathways. PLOS ONE. 2014;9(10):e110631.

42. Geller R, Taguwa S, and Frydman J. Broad action of Hsp90 as a host chaperone required for viral replication. Biochimica et biophysica acta. 2012;1823(3):698–706.

43. Amraiz D ZN, Fatima M. Antiviral evaluation of an Hsp90 inhibitor, gedunin, against dengue virus. Trop J Pharm Res. 2017;16(5):997–1004.

44. Vrijsen R, Vanden Berghe DA, Vlietinck AJ, and Boeyé A. Lycorine: a eukaryotic termination inhibitor? The Journal of biological chemistry. 1986;261(2):505–7.

45. Yang L, Zhang JH, Zhang XL, Lao GJ, Su GM, Wang L, et al. Tandem mass tag-based quantitative proteomic analysis of lycorine treatment in highly pathogenic avian influenza H5N1 virus infection. PeerJ. 2019;7:e7697-e.

46. Ieven M, van den Berghe DA, and Vlietinck AJ. Plant antiviral agents. IV. Influence of lycorine on growth pattern of three animal viruses. Planta medica. 1983;49(2):109–14.

47. Szlávik L, Gyuris A, Minárovits J, Forgo P, Molnár J, and Hohmann J. Alkaloids from Leucojum vernum and antiretroviral activity of Amaryllidaceae alkaloids. Planta medica. 2004;70(9):871–3.

48. Li SY, Chen C, Zhang HQ, Guo HY, Wang H, Wang L, et al. Identification of natural compounds with antiviral activities against SARS-associated coronavirus. Antiviral Res. 2005;67(1):18–23.

49. Liu J, Yang Y, Xu Y, Ma C, Qin C, and Zhang L. Lycorine reduces mortality of human enterovirus 71-infected mice by inhibiting virus replication. Virology journal. 2011;8:483.

50. Li S, Liu X, Chen X, and Bi L. Research Progress on Anti-Inflammatory Effects and Mechanisms of Alkaloids from Chinese Medical Herbs. Evidence-Based Complementary and Alternative Medicine. 2020;2020:1303524.

51. Rajapakse AG, Ming XF, Carvas JM, and Yang Z. The hexosamine biosynthesis inhibitor azaserine prevents endothelial inflammation and dysfunction under hyperglycemic condition through antioxidant effects. American journal of physiology Heart and circulatory physiology. 2009;296(3):H815–22.

52. Botz B, Bölcskei K, Kereskai L, Kovács M, Németh T, Szigeti K, et al. Differential regulatory role of pituitary adenylate cyclase-activating polypeptide in the serum-transfer arthritis model. Arthritis & rheumatology (Hoboken, NJ). 2014;66(10):2739–50.

53. Sasaki T, Kishi M, Saito M, Tanaka T, Higuchi N, Kominami E, et al. Inhibitory effect of di- and tripeptidyl aldehydes on calpains and cathepsins. Journal of enzyme inhibition. 1990;3(3):195–201.

54. Schneider M, Ackermann K, Stuart M, Wex C, Protzer U, Schätzl HM, et al. Severe acute respiratory syndrome coronavirus replication is severely impaired by MG132 due to proteasome-independent inhibition of M-calpain. J Virol. 2012;86(18):10112–22.

55. Upla P, Marjomäki V, Nissinen L, Nylund C, Waris M, Hyypiä T, et al. Calpain 1 and 2 are required for RNA replication of echovirus 1. J Virol. 2008;82(3):1581–90.

56. Zheng K, Xiang Y, Wang Q, Jin F, Chen M, Ma K, et al. Calcium-signal facilitates herpes simplex virus type 1 nuclear transport through slingshot 1 and calpain-1 activation. Virus research. 2014;188:32–7.

57. Ma C, Sacco MD, Hurst B, Townsend JA, Hu Y, Szeto T, et al. Boceprevir, GC-376, and calpain inhibitors II, XII inhibit SARS-CoV-2 viral replication by targeting the viral main protease. Cell research. 2020;30(8):678–92.

58. Barnard DL, Hubbard VD, Burton J, Smee DF, Morrey JD, Otto MJ, et al. Inhibition of severe acute respiratory syndrome-associated coronavirus (SARSCoV) by calpain inhibitors and beta-D-N4-hydroxycytidine. Antiviral chemistry & chemotherapy. 2004;15(1):15–22.

59. Blanc F, Furio L, Moisy D, Yen HL, Chignard M, Letavernier E, et al. Targeting host calpain proteases decreases influenza A virus infection. American journal of physiology Lung cellular and molecular physiology. 2016;310(7):L689–99.

60. Li X, Li Y, Shan L, Shen E, Chen R, and Peng T. Over-expression of calpastatin inhibits calpain activation and attenuates myocardial dysfunction during endotoxaemia. Cardiovascular research. 2009;83(1):72–9.

61. Supinski GS, and Callahan LA. Calpain activation contributes to endotoxin-induced diaphragmatic dysfunction. American journal of respiratory cell and molecular biology. 2010;42(1):80–7.

62. DeBiasi RL, Edelstein CL, Sherry B, and Tyler KL. Calpain inhibition protects against virus-induced apoptotic myocardial injury. J Virol. 2001;75(1):351–61.

63. Baker JD, Uhrich RL, Kraemer GC, Love JE, and Kraemer BC. A drug repurposing screen identifies hepatitis C antivirals as inhibitors of the SARS-CoV2 main protease. PLOS ONE. 2021;16(2):e0245962.

64. Sugamata R, Sugawara A, Nagao T, Suzuki K, Hirose T, Yamamoto K, et al. Leucomycin A3, a 16-membered macrolide antibiotic, inhibits influenza A virus infection and disease progression. The Journal of antibiotics. 2014;67(3):213–22.

65. Chiem K, Morales Vasquez D, Park J-G, Platt RN, Anderson T, Walter MR, et al. Generation and Characterization of Recombinant SARS-CoV-2 Expressing Reporter Genes. Journal of virology. 2021;95(7):e02209–20.

66. Chiem K, Ye C, and Martinez-Sobrido L. Generation of Recombinant SARS-CoV-2 Using a Bacterial Artificial Chromosome. Current protocols in microbiology. 2020;59(1):e126.

67. Sun C, Gardner CL, Watson AM, Ryman KD, and Klimstra WB. Stable, high-level expression of reporter proteins from improved alphavirus expression vectors to track replication and dissemination during encephalitic and arthritogenic disease. Journal of virology. 2014;88(4):2035–46.

68. Zou G, Xu HY, Qing M, Wang Q-Y, and Shi P-Y. Development and characterization of a stable luciferase dengue virus for high-throughput screening. Antiviral research. 2011;91(1):11–9.

69. Johannessen CM, Boehm JS, Kim SY, Thomas SR, Wardwell L, Johnson LA, et al. COT drives resistance to RAF inhibition through MAP kinase pathway reactivation. Nature. 2010;468(7326):968–72.

70. Ivens T, Eynde CVd, Acker KV, Nijs E, Dams G, Bettens E, et al. Development of a homogeneous screening assay for automated detection of antiviral agents active against severe acute respiratory syndrome-associated coronavirus. Journal of virological methods. 2005;129(1):56–63.

